# Selection-Enriched Genomic Loci (SEGL) Reveals Genetic Loci for Environmental Adaptation and Photosynthetic Productivity in *Chlamydomonas reinhardtii*

**DOI:** 10.1101/2021.07.06.451237

**Authors:** Ben F. Lucker, Nicolas L. Panchy, Joshua A. Temple, Urs F. Benning, Jacob D. Bibik, Peter G. Neofotis, Joseph C. Weissman, Ivan R. Baxter, Shin-Han Shiu, David M. Kramer

## Abstract

This work demonstrates an approach to produce and select hybrid algal strains exhibiting increased photosynthetic productivity under multiple environmental conditions. This simultaneously addresses two major impediments to improving algal bioenergy production: 1) generating new genetic variants with improved performance; and 2) disentangling complex interactions between genetic and physiological factors contributing to these improvements. We pooled progeny generated from mating two environmental isolates of the green alga *Chlamydomonas reinhardtii and* cultured the pools under multiple environmental conditions. Strains from the outcompeting populations showed substantial (in some cases over 3 fold) increases in productivity over the parental lines under certain environments related to biomass production, including laboratory conditions as well as hyperoxia, fluctuating light, high salinity and high temperature. The results indicate that *C. reinhardtii* has remarkable, untapped, directed evolution capacity that may be harnessed using breeding and competition approaches. The populations were deep sequenced at multiple time points to identify “Selection-Enriched Genomic Loci” (SEGL) that accumulated in the populations, and thus likely confer increased fitness under the respective environmental conditions. With improved resolution, SEGL mapping can identify allelic combinations used for targeted breeding approaches, generating elite algal lines with multiple desirable traits, as well as to further understand the genetic and mechanistic bases of photosynthetic productivity.

**Significance Statement:** Increasing the photosynthetic efficiency of algae during biomass production is perhaps the most critical hurdle for economically sustainable algal based biofuels. This presents unique challenges because modifications designed to increase photosynthesis often result in decreased fitness, due to production of toxic reactive oxygen species when photosynthesis is not adequately regulated. These problems are exacerbated under natural and outdoor production environments because of the complex nature of photosynthesis and the multifaceted interactions between genetic, environmental and physiological factors. Here, we demonstrate a high throughput biotechnological screening approach that simultaneously produces algal strains with highly increased autotrophic productivity and identifies genomic loci contributing to these improvements. Our results demonstrate that *Chlamydomonas reinhardtii* exhibits high directed evolutionary capacity readily accessed through breeding and selection.

## Background

A critical hurdle to achieving economically sustainable algal biofuel production is increasing the efficiency of photosynthetic growth in outdoor environments(1), in part because environmental factors in the real world decrease plant and algal photosynthetic efficiency far below theoretical maxima (2, 3). Further complications are that outdoor production conditions are subject to invasion by native algae, predators, and exposure to rapid changes in temperature and solar irradiance (4). Thus, meaningful improvements to algal productivity requires increasing photosynthetic efficiency while maintaining competitive fitness in exotic, dynamically changing environments.

Photosynthesis is a challenging trait to improve, in large part because it must be finely regulated to supply the correct amount and forms of energy to power cellular metabolic processes, while minimizing production of damaging Reactive Oxygen Species (ROS) that can occur when light energy input exceeds downstream sink requirements (5). To maintain viability, algae dissipate “excess” light energy under adverse environmental conditions (6–8), leading to fitness tradeoffs between efficient light capture and avoidance of photodamage. For example, decreasing photosynthetic antenna seize increases the photosynthetic efficiency of algae by optimizing the distribution of light through a culture (9–11). However, under natural or competitive conditions, small antenna phenotypes are negatively selected against due to reduced ability to compete for light resources (11, 12). A similar tradeoff is seen in *Chlamydomonas reinhardtii* (herein *Chlamydomonas*), where strains engineered to decrease energy loss through nonphotochemical quenching (NPQ) show increased growth under finely controlled conditions (13), but show reduced productivity and increased ROS accumulation under more production-like conditions (14), likely because increased light capture overwhelms the capacity of the background cellular physiology. Thus, photosynthetic improvements for outdoor cultivation requires consideration of complex interactions between genetic, environmental and physiological factors (6–8, 15), beyond targeted molecular biology efforts to adjust one process at a time.

In order for an alga to be grown as a monoculture for biomass production in outdoor environments, the strain will require a high degree of environmental fitness. It is perhaps only possible to achieve the level of fitness required for successful production through “domestication”. By definition, domestication occurs when a species is exposed to a selection environment associated with human cultivation and use (16). Humans are estimated to have semi- or fully domesticated 1,000-2,500 plant species exist from about 120-160 taxonomic families (16, 17). To date, 250 plant species are fully domesticated, with the overwhelming majority being angiosperms (18–20).

Clearly, in order for algae to become “domesticated” for biofuel or bioproduct targeted cultivation, domestication needs to become more rapid than the thousands of years it has taken for maize, wheat and other cereal crops (21–26). Fortunately, since the 1990s, the application of QTL mapping has demonstrated that, in many plants, we can begin to track the genetic basis of various domestications (27–33). A limited number of domestication experiments using artificial selection, have converted wild or crop-wild hybrids to domesticated forms in <20 generations (34–36). Similarly, evidence of severe genome-wide reductions in variation caused by bottlenecks and selection (37–41) and large selection sweeps (42–45) have suggested that domestication can occur rapidly. Together, it appears that domestication need not be a slow process, especially if strong selection pressure is applied to a breeding population (46).

By statistically associating genetic-linked phenotypic variations, it is possible to identify the genomic regions, or quantitative trait loci (QTL), contributing to increased vigor. Bulked segregant analyses (BSA) is a common QTL mapping approach, where segregant lines exhibiting extreme phenotypic differences are pooled and deep sequenced to find enabling genetic polymorphisms (47, 48). These, and other genome wide association approaches are powerful tools to identify the underlying locus, or combinations of loci, contributing to natural variations of complex traits (49–53). The potential of applying QTL-type mapping approaches for microalgae has been recognized (54–56), but not yet fully realized. *Chlamydomonas* is an ideal organism to explore these approaches because it is well-studied, genetically tractable, haploid and has a well-defined sexual cycle (57). Sexual crossing also plays a major role in the process of creating a domesticate, with interspecific hybridization having occurred in many crop species (21). In yeast, sexual recombination enables higher rates of adaptation compared to random mutagenesis,(58–62) suggesting that the sexual recombination capability of *Chlamydomonas* is a major asset.

Here, our strategy drew on several previously developed approaches, first crossing phenotypically distinct parent lines to generate segregant libraries similar to Recombinant Inbred Lines (RILs) used in many plant breeding studies (63). However, we adapted these approaches for photosynthetic microbes to simultaneously select for high efficiency and robustness, by pooling progeny and competing them under production-like conditions. By maintaining constant cell numbers under selection pressure, we enriched for robust lines that outcompete segregants with lower fitness or productivity. Individuals isolated from the outcompeting populations showed increased fitness under a wide range of selection conditions, including high oxygen, high salt, prolonged light deprivation, alternating high and low light and elevated temperature. Many of these lines show little tradeoffs under non-selection conditions, suggesting that *Chlamydomonas* has strong, untapped directed evolutionary capacity that could be exploited to generate highly efficient production lines. Using a new statistical approach that can account for selection-induced changes in the genomic population, we identified Selection-Enriched Genomic Loci (SEGL) that are likely to contribute to the increased vigor. Our results 1) demonstrate that Chlamydomonas can be utilized as a platform to map the loci which may increase the organism’s productivity in a given environment relevant to biofuel production and 2) our methods can be used to breed an algae better suited for a specific condition, thus initiating the domestication of an algae, particularly, Chlamydomonas.

## Results

### Allele frequency tracking in mixed populations by SNP mapping

We screened a series of natural isolates and laboratory strains of *Chlamydomonas* for productivity under three well-defined conditions: (1) baseline condition that mimics a natural solar day (BC, 5% CO_2_ in air, 14:10 light dark cycle with zenith at noontime); (2) hyperoxia (HO), as is often encountered under mass production conditions(64) (5% CO_2_ in O_2_, other conditions as in BC); and (3) light stress (LS), (BC days followed by three days of very low light), to favor lines that can rapidly store fixed carbon on “light replete days”, and maintain growth during the “light starvation” days (see Materials and Methods and Fig. S1). One pair of isolates, CC1009 and CC2343, exhibited similar growth under BC, but substantially different growth under HO and LS conditions (Fig. 1A). Under LS, CC1009 reduced productivity by 20% during the light replete days, but CC2343 growth was essentially completely inhibited. Neither line was able to grow on the light starvation days. Under HO conditions, CC1009 lost 66% of productivity whereas CC2343 lost 87%.

**Figure 1.**
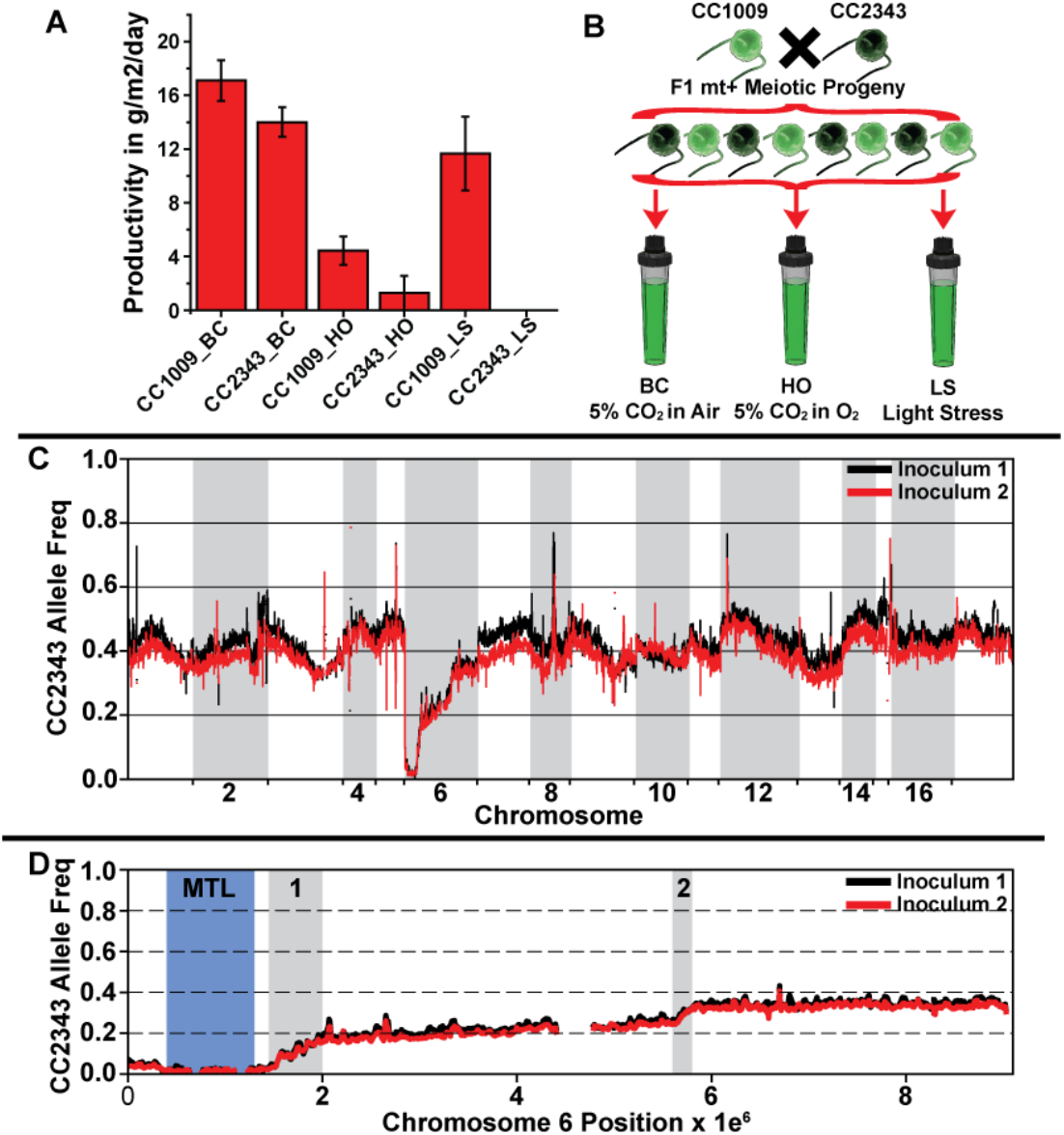
The experimental approach used to generate and map diverse populations. (A) The average daily biomass productivity of the selected parental strains (CC1009 and CC2343) under BC (n=8), HO (n=5), and LS (n=5) conditions. Error bars represent the standard deviation between biological replicates. (B) The F1 competition experimental outline. Parental lines CC1009 and CC2343 were crossed and 203 F1 mt- progeny were pooled in equal numbers and used as inoculum for cultures placed under either BC, HO or LS conditions. (C) The allele frequency relative to CC2343 in 40 Kb windows across the genome for two independently generated inoculums, BC and HO experiment (Inoculum 1, black line), BC and LS experiment (inoculum 2, red line). (D) The allele frequency relative to CC2343 across chromosome 6 for the inoculums in panel B. The blue shaded regions highlights the mating type locus (MTL) while the grey shaded regions are potential regions for increased recombination rates.

To generate a diversity panel of progeny, we crossed CC1009 (mt-) with CC2343 (mt+) (Fig. 1B). Using a refined list of Single Nucleotide Polymorphisms (SNPs) identified from the parental strains (see SI Materials and Methods), we mapped the CC2343 allele frequency of two independently generated inoculum populations used for competition experiments (Fig. 1C). The similar allele frequency distributions across the genomes of the F1 inoculums show that population pooling, deep sequencing and methods to quantify the frequencies of SNPs generated reproducible results. Excluding chromosome 6, the CC2343 allele frequency varied between 0.5 and 0.35 across the genome. Averaging all of the allele frequencies across the genome generated frequencies of 0.42 and 0.58 for CC2343 and CC1009 respectively, indicating a slight bias for CC1009 loci within the population. By contrast, the 700 kb segment of DNA at the beginning of chromosome 6, corresponding to the mating type locus(65, 66), showed strong selection for CC1009 loci. This served as an internal positive control, because we exclusively selected mt- strains for the F1 competition experiments, so that the population should be essentially homozygous for the CC1009 mt- mating type locus, while nearby (linked) loci should strongly favor CC1009 (Fig. 1D, blue shaded area). CC1009 loci was progressively lost moving away from the mating type locus, indicating that crossover events must have occurred following mating. The largest changes of allele frequency on chromosome 6 occurred in two distinct regions, together totaling <1 MB (Fig. 1D, grey shaded regions), suggesting the possibility of recombination hotspots.

### Competitive growth under different conditions selects for distinct combinations of genomic loci

Pooled F1 inoculums were cultured in environmental photobioreactors (67) and grown under either BC, HO or LS conditions, and samples were collected periodically for deep sequencing during each competition (see Table S1 and Fig. 1B for replicates and sequencing coverage). The changes in allele frequencies for each population were rapid and unique for each population (Figs. 2 a, c, e, S2-S5), and highly statistically significant as shown by the LOD (log of odds) scores (described in Materials and Methods), (Figs. 2 b, d, f). Biological replicates under each condition showed very similar patterns and extents of allele frequency changes (Figs S2-S5), and high correlations between replicates (Fig S6).

**Figure 2.**
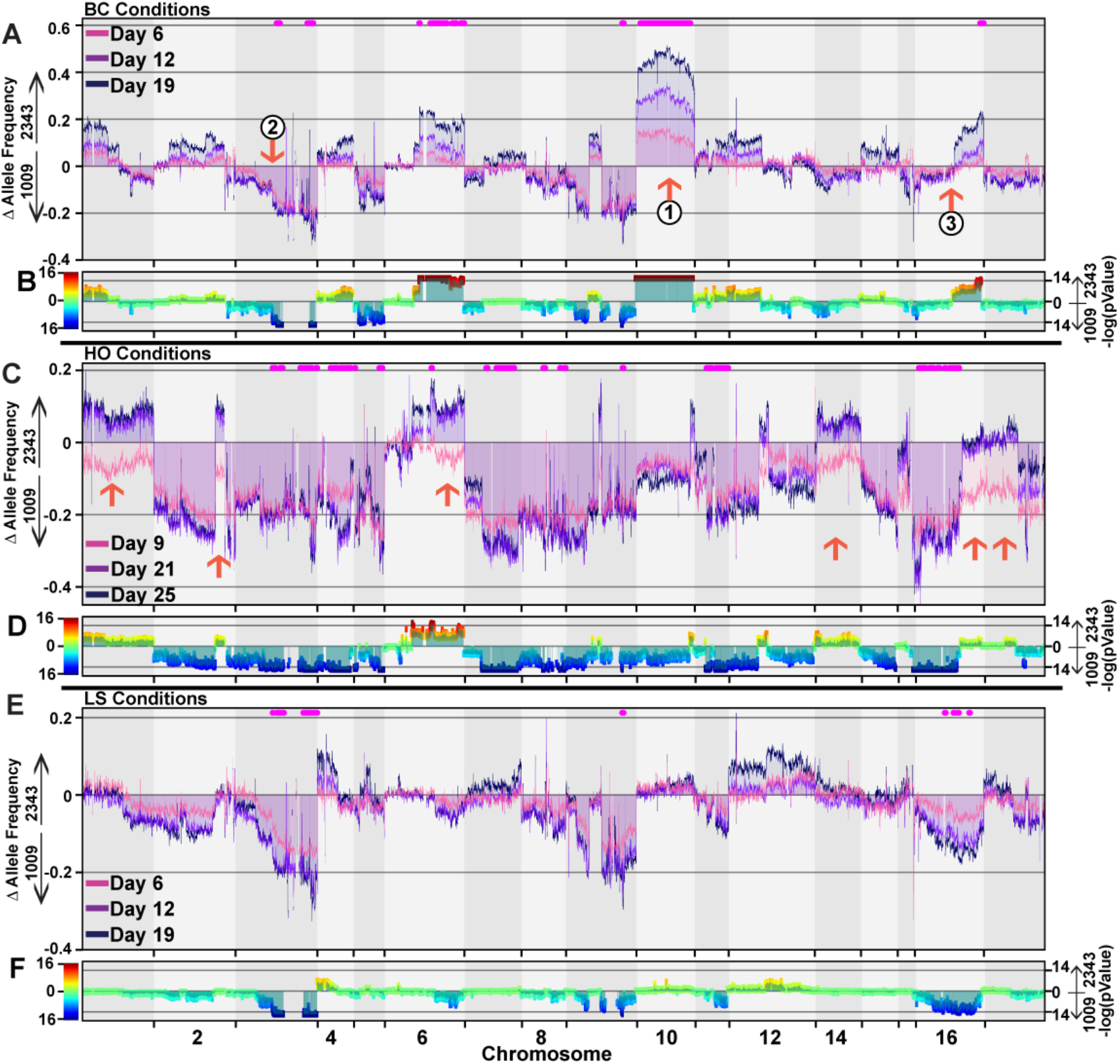
Environmental conditions drive population genome structure. (A) The change in the allele frequency of the F1 BC population from the inoculum relative to CC2343 (positive numbers) or CC1009 (negative numbers) after 6 (violet), 12 (magenta), 19 (dark blue) days in polyculture. (B) The degrees of allelic enrichment (-log_10_(p)) of 40 Kb windows with a step size of 8 Kb across the genome for the BC population after 19 days of polyculture (dataset from the F1 LS experiment). As described in SI Materials and Methods, *p* is the probability of achieving the observed allele frequency change of a loci randomly. Regions with -log10(p)>14 (a highly stringent cutoff) over a 60 kb window were considered to be an Enriched Genomic Locus (SEGL). To illustrate the relative preference for loci from the two parents, - log10(p) values of SEGL regions that represent enrichment of CC1009 were multiplied by -1. (C) The change in allele frequency of the F1 HO population after 9 (violet), 12 (magenta), 19 (dark blue) days in polyculture. (D) The enrichment values from the F1 HO population after 21 days of polyculture. (E) shows the change in allele frequency of the F1 LS population after 6 (violet), 12 (magenta), 19 (dark blue) days in polyculture. (F) the enrichment values from the F1 LS population after 19 days of polyculture.

Despite the fact that BC was designed as a low stress condition, it imposed strong selection for specific loci throughout the genome (Figs. 2A, S2, S4). However, the aggregated frequency of parental alleles across the genome showed only a slight (∼2.3%) preference for CC2343 at the end of the competition (Table S2). Competitions under “harsher” conditions of HO and LS resulted in stronger enrichment for alleles from CC1009 over CC2343, and the final populations showed 15% and 3% increases in CC1009 alleles respectively (Table S2). These results likely reflect selection for hybrid progeny containing alleles that confer higher stress tolerance related to CC1009. Nonetheless, some CC2343 alleles were enriched as well, indicating novel genetic combination from both strains contributed to the fitness of the final population.

The kinetics (or time-dependencies) of allele frequency changes within the populations followed distinct patterns for different regions and conditions. Some regions showed gradual increases throughout the experiment (e.g. chromosome 10 under BC, see Fig. 2A, arrow 1), whereas chromosome 3 showed rapid initial selection in all conditions (e.g. Fig. 2A, arrow 2). In some cases, only part of the chromosome was selected for, while other regions remained unchanged (e.g. Fig. 2A, arrow 3), and in other cases, an initial selection for one loci was reversed at later times (Fig. 2C, arrows B). We interpret these complex behaviors as reflecting the importance of the most impactful loci first, followed by selection for secondary effects of allelic combinations that sometimes can be maladaptive. Under LS, the overall rates of changes were slower than under BC or HO, most likely due to reduced cell divisions during the low light days (see Table S2).

Comparing the variation in allele frequencies across our biological replicates, we calculated the LOD scores for each 60bp genomic windows to identify SEGLs (see Material and Methods). Using a highly conservative threshold of LOD>14, we identified 11, 12 and 52 SEGLs for the LS, BC and HO conditions, respectively. The SEGLs ranged in size between 64 Kb and the entirety of chromosome 10 at 6.5 Mb for BC conditions, but the median size of all identified SEGLs was 144 Kb. Some mapped SEGL were common between all conditions, such as the far-right side of chromosome 3 and a 140 Kb region of chromosome 9. However, each environmental condition led to selection for specific allelic combinations in each population (Figs. 2 and S2-S5). For example, on chromosome 16 BC selected for an SEGL of CC2343 alleles on the far-right end of the chromosome, while the HO SEGL on the left ∼60% of the same chromosome was for CC1009 loci, with the highest selective pressure on the far left. Contrasting this was the LS SEGL in the central region of chromosome 16 that was enriched for CC1009 loci. More specific examples can be observed by scanning the plots in Fig. 2.

We should note however, most chromosomes can be divided up into multiple sub-regions with substantial allele frequency changes, reminiscent of haplotype blocks. For example, on chromosome 6 of the HO population (at day 21), there were at least six sub-regions with allele frequency changes of approximately 0, -0.11, -0.02, 0.10, 0.18, and 0.10 from left to right (Fig. 2C). These sub-regions show significant differences in LOD scores (> 3) from their neighboring regions and often show distinct accumulation kinetics over time between environments (e.g. Fig. 2C, arrows), supporting the view that they represent distinct allelic combinations that confer differentially to selection. While it is tempting to suggest that distinct loci affecting fitness fall under these regions, additional resolution is needed to further break down large SEGL regions to narrowing down contributing genes.

### Mating-induced genomic diversity following an F1 cross and the effects on selection and SEGL mapping resolution

To better understand the impact of increased population size and cross-over events that contribute to distinct allelic combinations on our SEGL mapping potential, we generated an F2 population of >240 lines by intercrossing F1 progeny (Fig. 3A) from two dissected tetrads. The F1 progeny used to generate the F2 lines were sequenced and contributing CC1009 and CC2343 loci were mapped (Figs. 3B and S7). The F1 progeny showed an average of about 13 crossover events for each cell, distributed over the 17 chromosomes, giving us an estimate for the rate of genetic diversification during meiosis in *Chlamydomonas*. There were multiple crossover events at common loci between the two meiotic events on chromosomes 9, 13, 14, and 17 (Fig. S7), suggesting again the possibility of recombination hotspots as observed in many multicellular plant species(68). Surprisingly, the distribution of CC1009 and CC2343 allele frequencies in the pooled F2 inoculum deviated from the expectation of equal contribution from each of the F1 parents (Fig. 3D). These deviations showed transitions at distinct points on the genome, many of which coincided with crossover locations in the contributing F1 lines, e.g. compare arrows in Fig. 3C for transitions on chromosome 2, with specific crossover events in the F1 lines (arrows in Fig. 3B). These effects suggest that the second mating itself led to selection for certain genomic loci, perhaps because of genomic incompatibility causing loss of viability, limiting the overall diversity of the population. It is also possible that at the time of the library pooling some lines had gone through S-phase and contained twice the number of chromosomes. In any case, the results show that one cannot assume complete segregation of parental loci, and it is critical to compare enrichment results to the initial allele frequency distributions.

**Figure 3.**
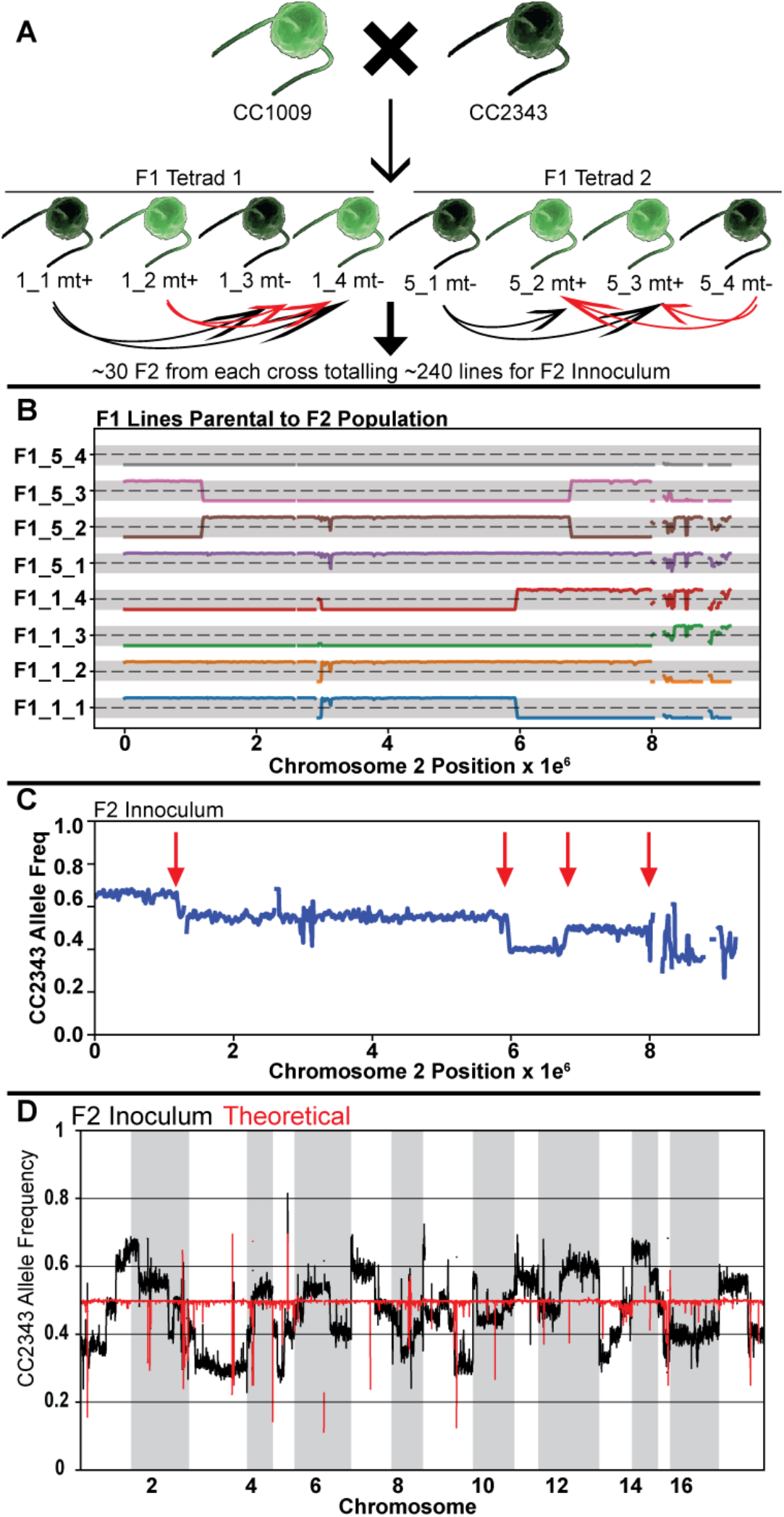
F1 recombination events shape F2 population genome structure. (A) Shows the breeding paradigm to generate the F2 progeny library. Two F1 tetrads were dissected and crossed with the two opposite mating type from the same tetrad and 30 F2 progeny from each cross were pooled to generate the F2 progeny library of ∼240 lines. (B) Shows the offset allele frequencies relative to CC2343 of chromosome 2 for the dissected tretrad progeny, which are the F1 progeny used to generate the F2 population. The allele frequencies range from 0 to 1 and are centered on the horizontal dotted lines at a relative allele frequency of 0.5. (C) Shows the chromosome 2 allele frequency of 240 pooled F2 lines used as the F2 inoculum. (D) The allele frequency across the genome of the F2 inoculum (black line) deviates from theoretical values (red line). The theoretical allele frequency was calculated by averaging the allele frequency from each F1 progeny cross shown in panel A.

**Figure 4.**
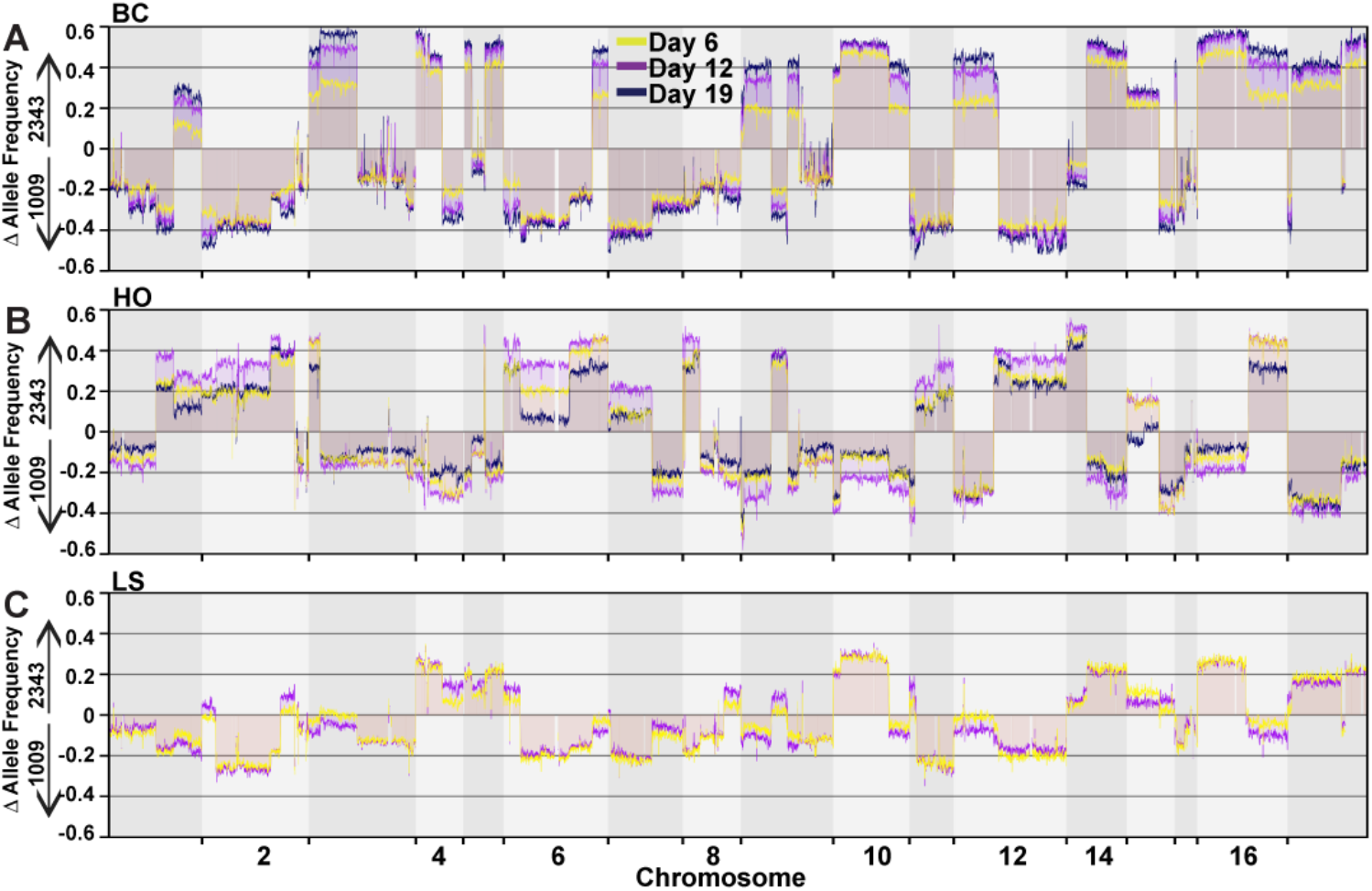
F2 populations show a bimodal distribution of progenitor loci. The F2 library was used to inoculate triplicate ePBRs and the cultures were placed under BC, HO, and LS conditions. (A) Shows the average change of the allele frequency of the F2 BC population after 8 (cream), 16 (violet), and 21 days (dark blue). (B) Shows the average allele frequency change of the F2 HO population for the same time points as panel A. (C) Shows the average change in allele frequency for the F2 LS population after 8 (violet) and 16 days (dark blue).

The F2 population (Fig. 3D) was competed under BC, HO and LS conditions and samples were collected at days 8, 16 and 21 for deep sequencing to track the allele frequency of each population (Tables S1 and S2). The allele frequencies between replicate populations were highly reproducible for all three conditions (Fig. S8-S10) and showed high degrees of correlation (Fig S11). As with the F1 pool, competing the F2 population under different environmental conditions led to enrichment of distinct combinations of genomic regions, but with some important differences. The F1 competitions resulted in approximately Gaussian distributions of allele frequencies, implying that the final pool contained a range of genetic variants that were able to compete relatively evenly. By contrast, the F2 competition, particularly under BC, led to selection for regions from mostly one progenitor or the other (Fig. S12) resulting in a bimodal distribution of allele frequencies, preventing us from obtaining SEGL mapping of the F2 population (see Fig. S8, day 21). The bimodal allele frequency distributions (Figs. S8-10, S12) are consistent with a strong founder effect and that the competitive advantage for individuals in the F2 populations was likely resulting from specific allelic combinations inherited from the F1 parental lines. Supporting this interpretation, the F2 competed populations retained stretches of the chromosome that contains the crossover positions from the individual dissected F1 tetrads (Fig. S13).

Interestingly, each environmental condition selected for different genomic regions from the F1 parents. For example, BC selected almost exclusively for genomic regions matching crossovers in a single F1 tetrad (termed F1_5_4, see Fig. S13 panel A). To a lesser extent HO selected for loci from a different tetrad parent (F1_1_2) (see, e.g. see arrows in Figs S13 panels A and B indicating abrupt changes in allele frequency in chromosomes 1, 3, 7, 9, 11,13, and 16). LS produced a population with the highest genomic diversity, as shown by the more Gaussian distributions of allele frequency (Fig. S11 Panel C), and clear contributions from at least two tetrad parents (F1_5_3 and F1_5_4).

### Chlamydomonas shows strong evolutionary capacity

For breeding to effectively improve industrial algal production, it is critical that some meiotic progeny are more productive than their parent lines. Indeed, the growth rates (doublings day^-1^) of the competing, co-cultured F1 and F2 populations under BC and HO eventually surpassed that of either of the parental lines under the same conditions, suggesting that algal fitness was a transgressive trait and some of the outcompeting progeny possessed novel allelic combinations of different loci (Fig. S14). We then screened a subset of outcompeting lines, isolated at the end of F1 and F2 competitions. Strikingly, the majority of winners from both F1 and F2 populations showed higher productivity (or tolerance) than either of the original parent lines, under the respective competition conditions (Figs. 5A-C, S15-S16). The best performing winners from the F1 competition under BC and HO showed 20% and 210% increase biomass productivities, respectively, compared to the more productive progenitor line. Similar trends were observed with selected F2 winners that had varied degrees of improvement (see Figs. S15 and S16). These results indicate that the new combination of genes, shuffled from both parents led to individuals with more optimal phenotypes, and strains exhibiting these traits underwent positive selection.

**Figure 5.**
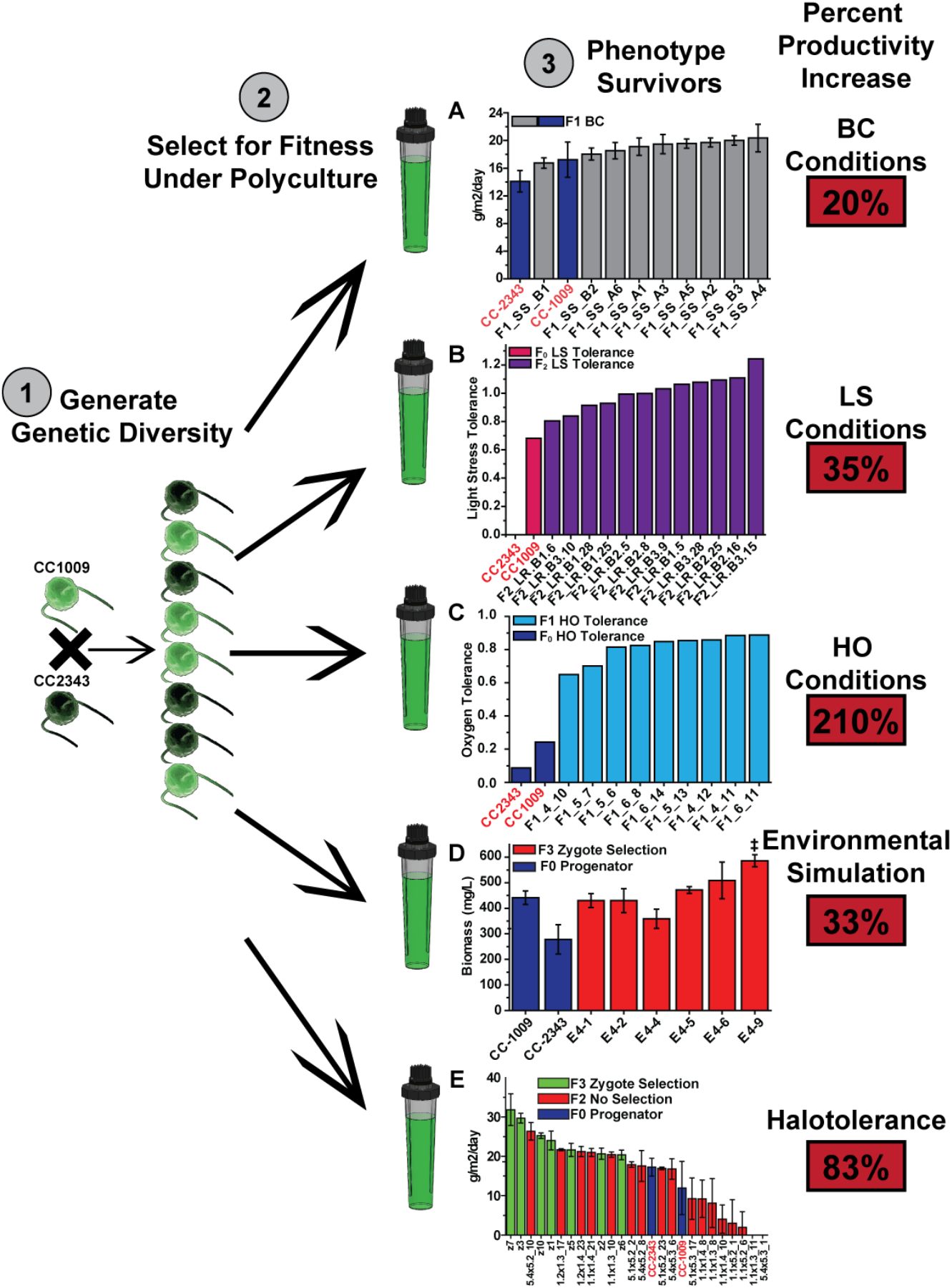
Through breeding and selection, *Chlamydomonas* shows high degrees of phenotypic plasticity. Step 1 is to generate genetic diversity through breeding divergent lines. Step 2 is to compete the lines under polyculture conditions. Step 3 is to isolate and screen the surviving progeny for increased productivity. (A) Shows average daily productivities of isolates from the F1_BC population (grey bars) and the parental strains CC1009 and CC2343 (dark blue bars for all panels). (B) The LS tolerance of surviving isolates of the F2_LS population (pink bars), is shown. (C) Shows the HO tolerance of F1_HO survivors (light blue bars) compared to the parental strains. (D) The productivity of selected progeny (red bars) compared to the parental strains after an environmental simulation is shown. Error bars represent the standard deviation of (from left to right) n = 4, 5, 5, 5, 4, 4, 4, 6 biological replicates (E) Summarizes a screen of progeny productivity in media containing 20 g/L of Instant Ocean. The green bars are strains isolated after hatching and selection under 20 g/L of Instant Ocean salts, red bars are random F2 progeny and blue bars are CC2343 and CC1009. Error bars represent the standard deviation between daily average productivities. It should be noted that the observed percent productivity increases under selection conditions are not equal to tolerance increases.

We also tested for tradeoffs in performance after comparing outcompeting lines selected under one condition on other conditions. Lines selected under HO also showed similar or even more robust growth under BC conditions (Figs. S15 B and E), suggesting that HO tolerance is not a tradeoff for BC tolerance. Also, nearly all lines selected under LS showed increased tolerance (a higher ratio of LS:BC productivity) and productivity under LS conditions, but all had decreased fitness under BC (Fig. S15 G, H). Overall, these results suggest that, while some traits selected for in our competitions may incur tradeoffs under other conditions, others appear to be compatible across multiple conditions relevant to the autotrophic mass production of algae.

To further explore the evolutionary capacity of Chlamydomonas, we streamlined our mating and selection process by hatching pools of hundreds to thousands of isolated F3 zygotes during, or just prior to the exposure to a new set of selection conditions. In the first experiment pooled zygotes were hatched prior to exposure to harsh pond-like conditions, with fluctuating temperatures (between 12 and 44 ^○^C) and high light (Fig. S16). Under these conditions, all of the selected winners performed better than the poor performing parent line, CC2343, and one showed a statistically significant increase (∼33%) over the better performing parent, CC1009 (Fig. 5D).

In a follow-up experiment, F3 zygotes were hatched directly under high salt conditions (same as BC but with temperatures held at 28 ^○^C and 20 g/L of Instant Ocean salts, see Materials and Methods) followed by culturing in turbidostat mode under these conditions for eight days prior to isolating lines from the outcompeting population. Seven randomly selected outcompeting lines showed increased productivity under high salt conditions ranging from between 20% and 83% compared to the more productive of the progenitor lines (CC2343) (Fig. 5E). For comparison, we also randomly selected seventeen F2 progeny prior to selection and grew them under high salt conditions, and found growth rates from zero (i.e. lethal) to 45% above that of the parent lines (Fig. 5E). This experiment clearly demonstrates that polyculture competition pre-screens strains for increased fitness prior to monoculture phenotyping. Taken together, these data demonstrate that through breeding and selection of *Chlamydomonas,* segregants can be generated which have increased fitness, with traits that allow higher productivity under simulated production conditions, including tolerance for high temperatures, salinity, and high oxygen conditions.

## Conclusions

We demonstrate that breeding natural variants of *Chlamydomonas* and selection under polyculture conditions can be used for streamlining enrichment for segregants showing higher growth and fitness under a wide range of environmental challenges. We also show that quantitative genomics approaches, such as BSA or SEGL, can be used to identify genomic loci that reflect the genetic bases for the observed increased fitness. In this study the resolution of SEGL regions in the F1 population spans regions from 60 KB to over 1.2 MB, encoding from 10 to over 2000 genes. Thus, although the resolution was too low to identify specific genes linked to increased productivity, results on both the F1 and F2 competitions suggest that increased SEGL resolution can be obtained by generating larger libraries of primary and secondary crossover events, followed by environmental selection.

Though we cannot dismiss that only a few loci are responsible for the observed improvements in productivity of the winners under any condition, the number of genetic loci, or quantitative trait loci (QTL) that underlay even a single trait related to domestication in other species has been often difficult to estimate (20). Even for single traits in maize, the number of QTL range from 6 to 26 (27). One study suggested that the nearly 500 genomic regions, spanning an estimated 2,000 genes contributed to domestication of maize (69). This suggests that, like many crop plants (70), the highest levels of improvement in algal photosynthesis may require stacking of many genetic variations with small effects. For *Chlamydomonas*, a library of at least 12 natural isolates and 25 lab strains have been sequenced, and some phenotypic analysis of the strains revealed substantial phenotypic variation between isolates and lab reared strains (54, 71–73), indicating that there is significant genetic diversity to drive artificial breeding efforts. It has been shown in *Chlamydomonas* that selection under mixotrophic or high light conditions can lead to higher fitness after mutagenesis (74–76) as well as higher levels of halotolerance after sexual recombination and selection (77–79). Thus, it appears that modern genomics-targeted breeding and phenotypic selection can form the basis for rapid improvement of photosynthetic productivity and identification of the enabling allelic combinations in order to domesticate an algae.

Crossing- and selection-induced gains in productivity observed under one condition did not necessarily impose tradeoffs of decreased productivities, and in some cases appeared to increase productivity under other conditions, suggesting that the approach may be used to stack multiple productivity and fitness traits for a specific set of environments. The system is highly scalable through high-throughput variant generation and bulk selection, and thus the approach may be especially relevant for generating production strains without the requirements for marker genes as in the case in conventional genetic engineering. Future analyses of the progeny with various performance levels in comparison to the parents should yield insights into the complex nature of how suites of genetic components and associated encoded proteins are coordinated to increase photosynthetic productivity and environmental fitness under specific conditions. These should be highly relevant to algae domestication for biomass production for biofuels and bioproducts.

## Acknowledgments

The Authors would like to thank Amy Terauchi and Jennifer Barrett (supported by USDS-ARS), Jonathan Flowers, and Barbara Sears for early intellectual contributions to this work. The initial proof of concept for the competition experiments, the subsequent work related to hyperoxia, and development of software for data analyses were supported by the US Department of Energy (DOE), Office of Science, Basic Energy Sciences (BES) award number DE-FG02-91ER20021. The analyses under multiple environmental conditions, development of statistical analyses and trade off assessments were supported by the ExxonMobil corporation. The development of the photobioreactor systems was supported by the US Department of Energy National Alliance for Algal Biofuels and Bioproducts (NAABB). Salary for DMK was partially funded by MSU AgBioResearch.

---

**Figure S1.**
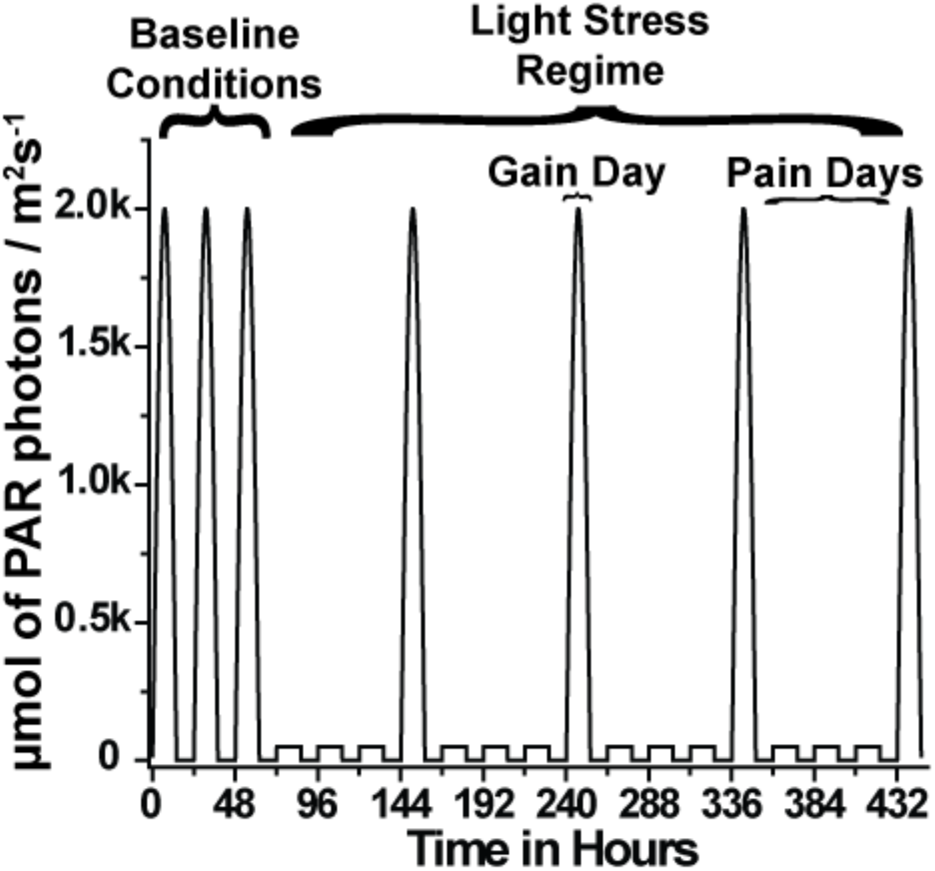
The culture light intensity in µmol of photosynthetically active radiation (PAR) photons per m^2^s^-1^ during baseline conditions (BC) and during the light stress regime (LS). It also defines “gain days” and “pain days”.

**Figure S2.**
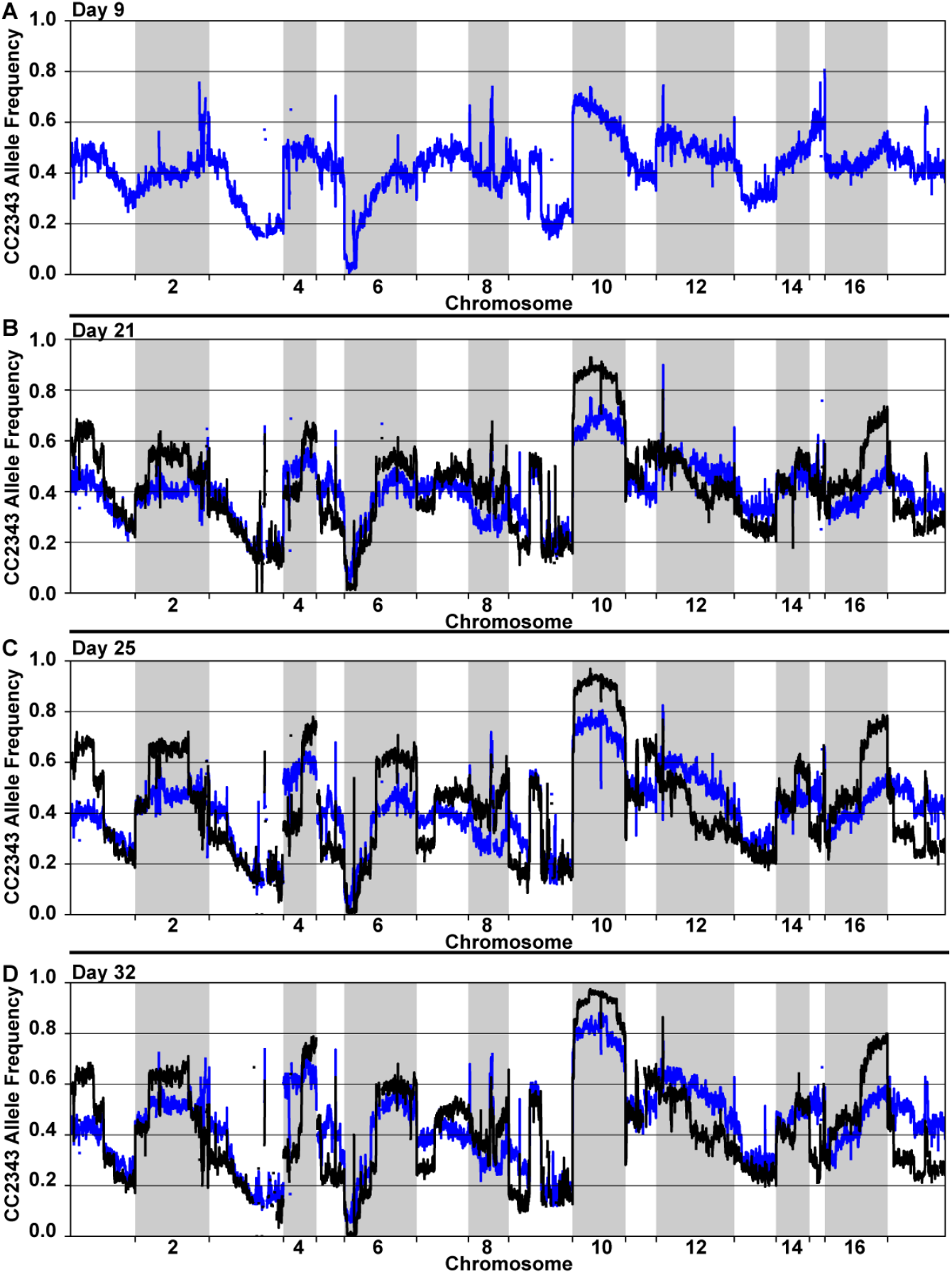
The allele frequency relative to CC2343 for replicate F1 BC populations at days 9 (A), 21 (B), 25 (C), and 32 (D) are shown. Allele frequencies were averaged over 40 Kb windows sliding every 8 Kb. Line colors represent individual replicates. Parallel experiments were performed on the F1 HO population using the same inoculum population.

**Figure S3.**
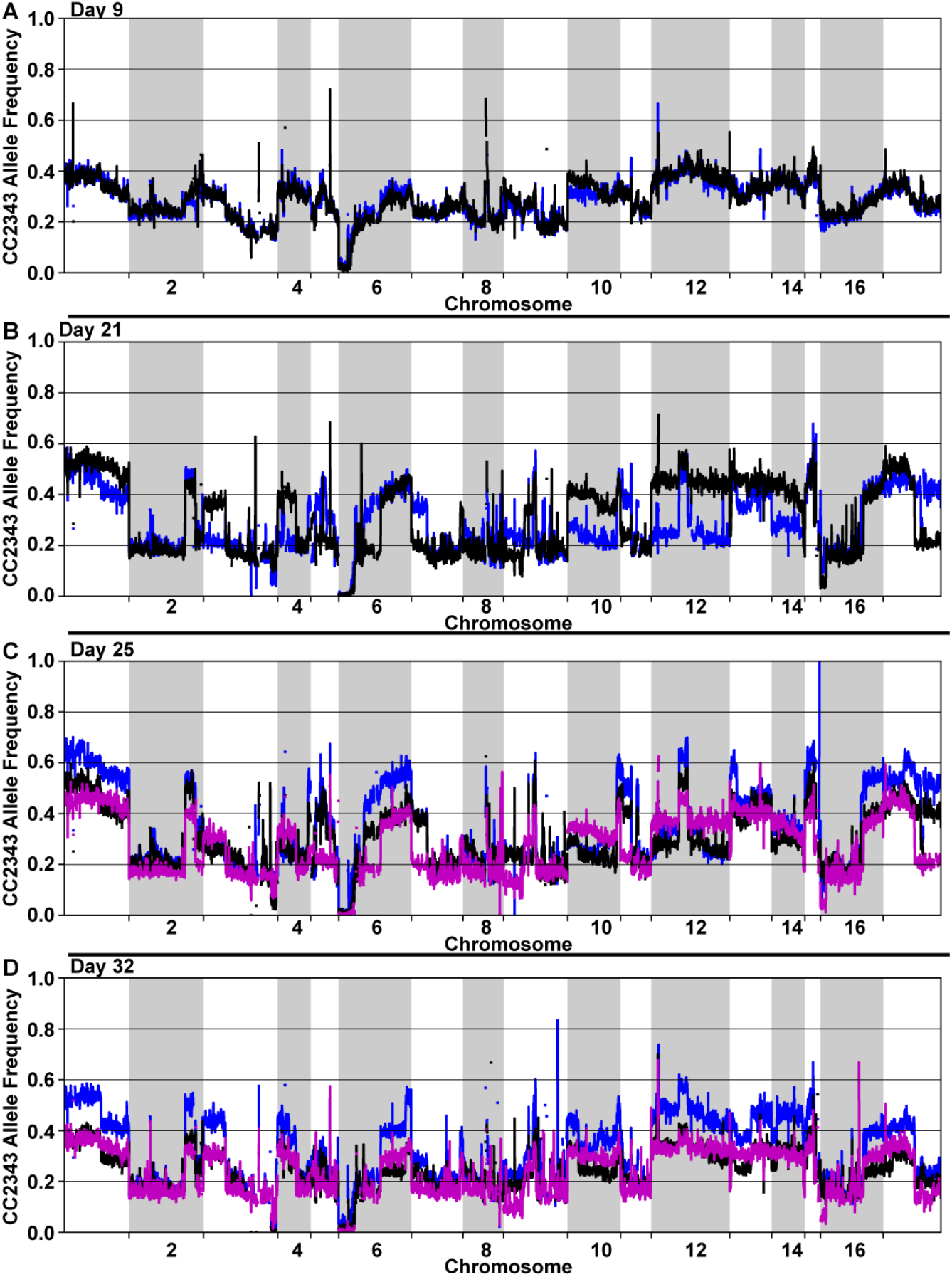
Allele frequencies relative to CC2343 for replicate F1 HO populations at days 9 (A), 21 (B), 25 (C), and 32 (D). Allele frequencies were averaged over 40 Kb windows sliding every 8 Kb. Line colors represent individual replicates.

**Figure S4.**
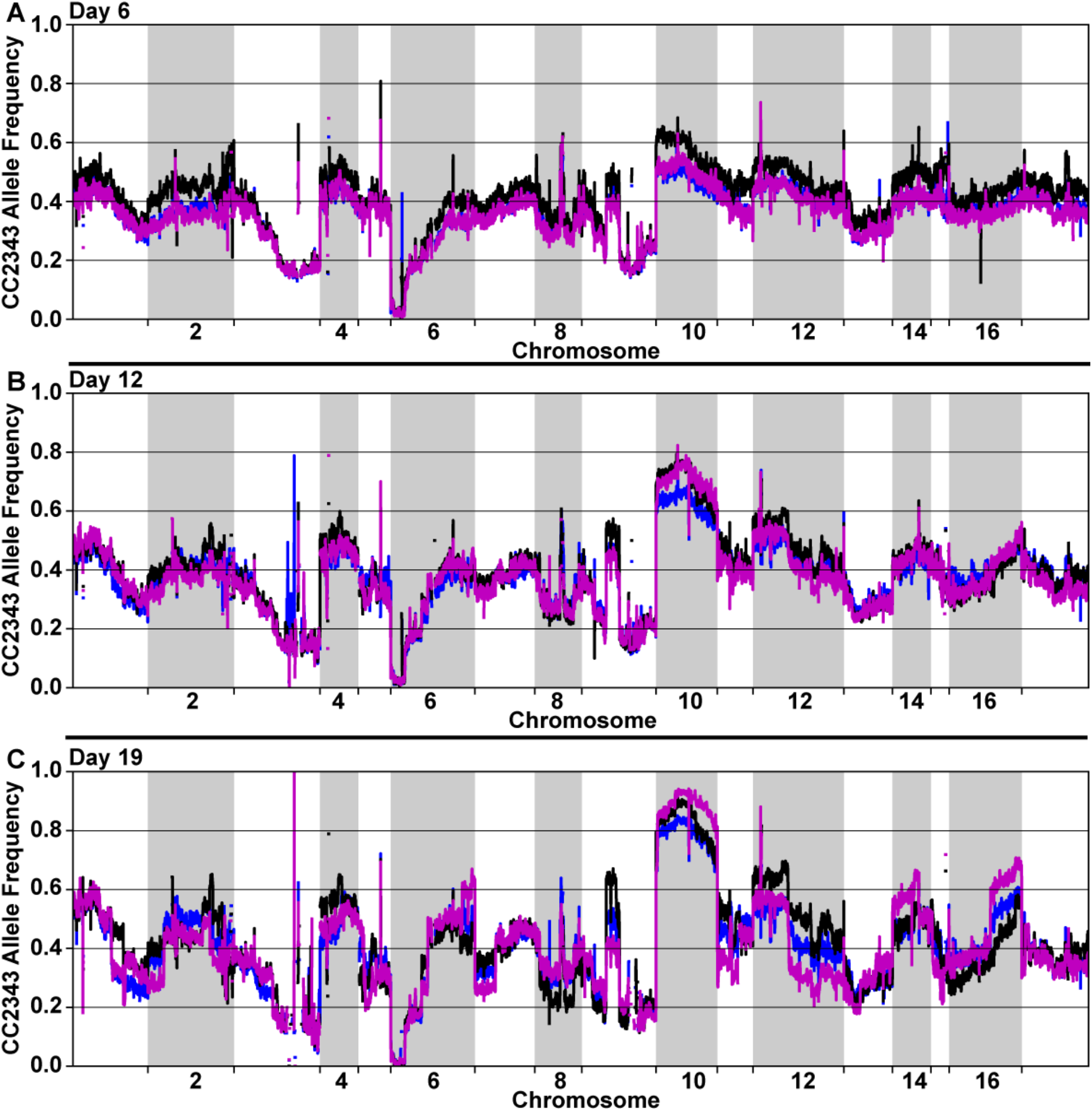
Allele frequency relative to CC2343 for replicate F1 BC populations at days 6 (A), 12 (B) and 19 (C). Allele frequencies were averaged over 40 Kb windows sliding every 8 Kb. Line colors represent individual replicates. Parallel experiments were performed with the F1 LS populations using the same inoculum.

**Figure S5.**
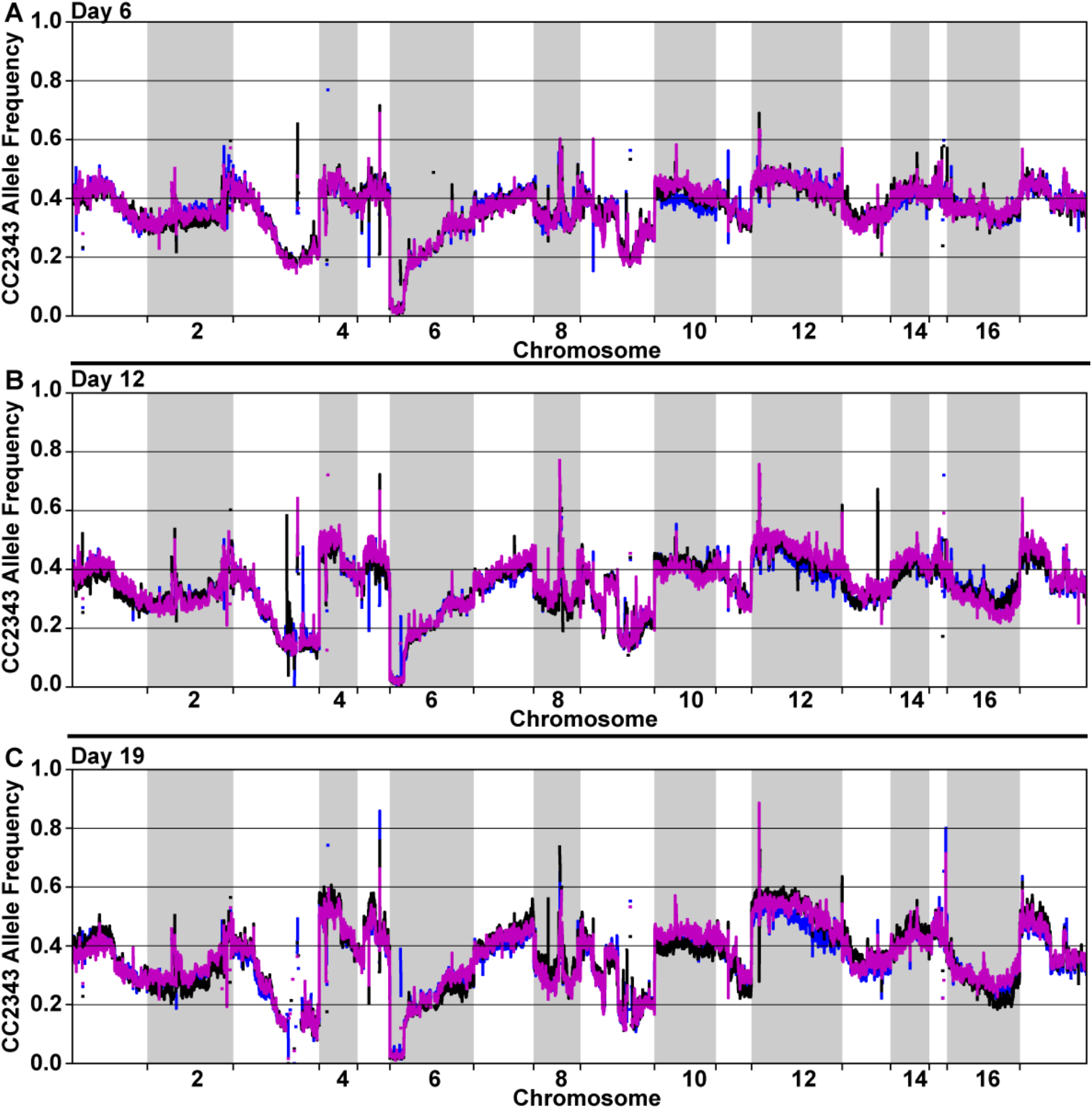
Shows the allele frequency relative to CC2343 for individual F1 LS populations at days 6 (A), 12 (B), and 19 (C). Allele frequencies were averaged over 40 Kb windows sliding every 8 Kb. Line colors represent individual replicates.

**Figure S6.**
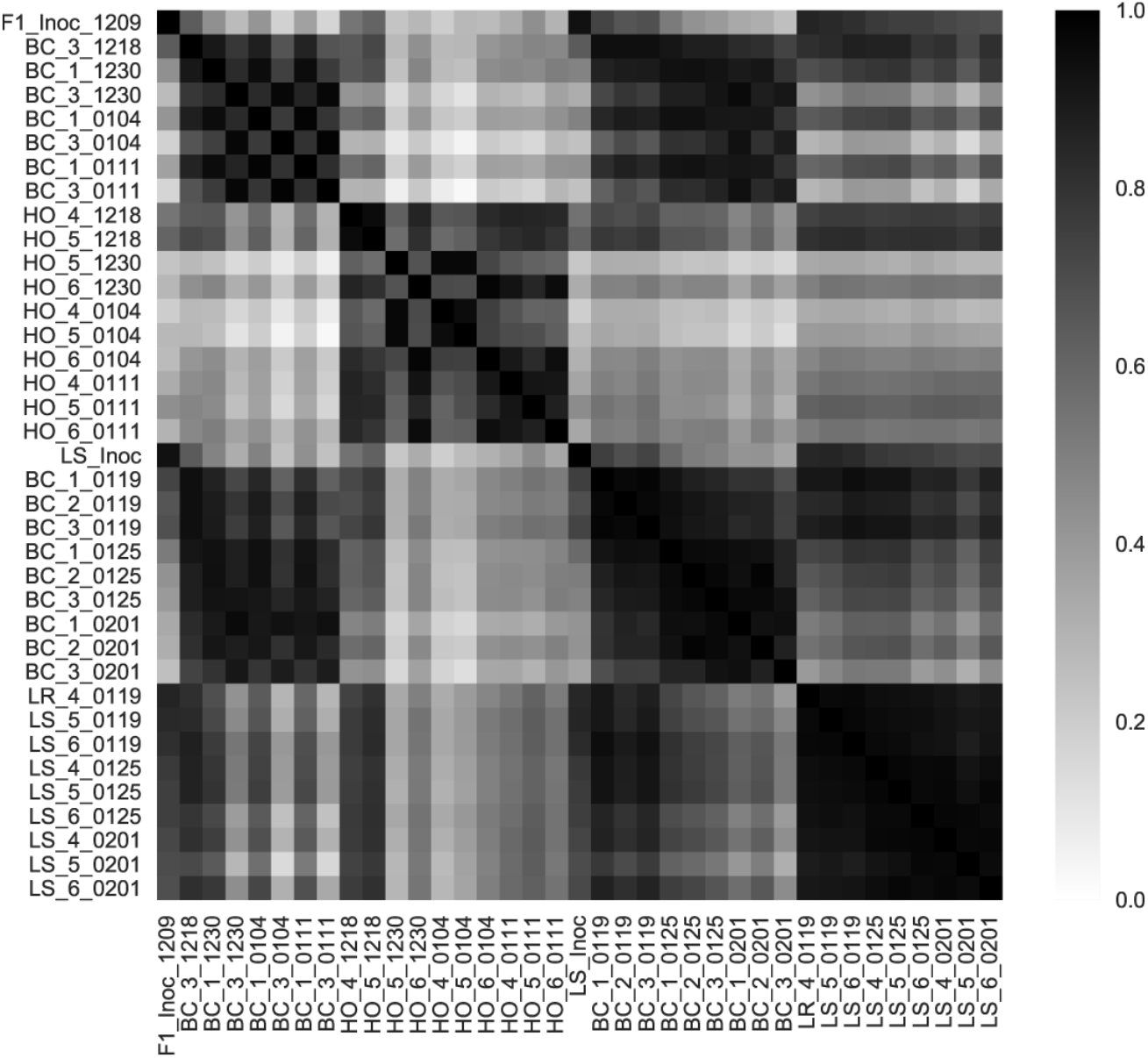
A high degree of reproducibility is shown using a matrix of Pierson Correlation Coefficients between the allele frequencies of replicate F1 populations exposed to the same environmental selection.

**Figure S7.**
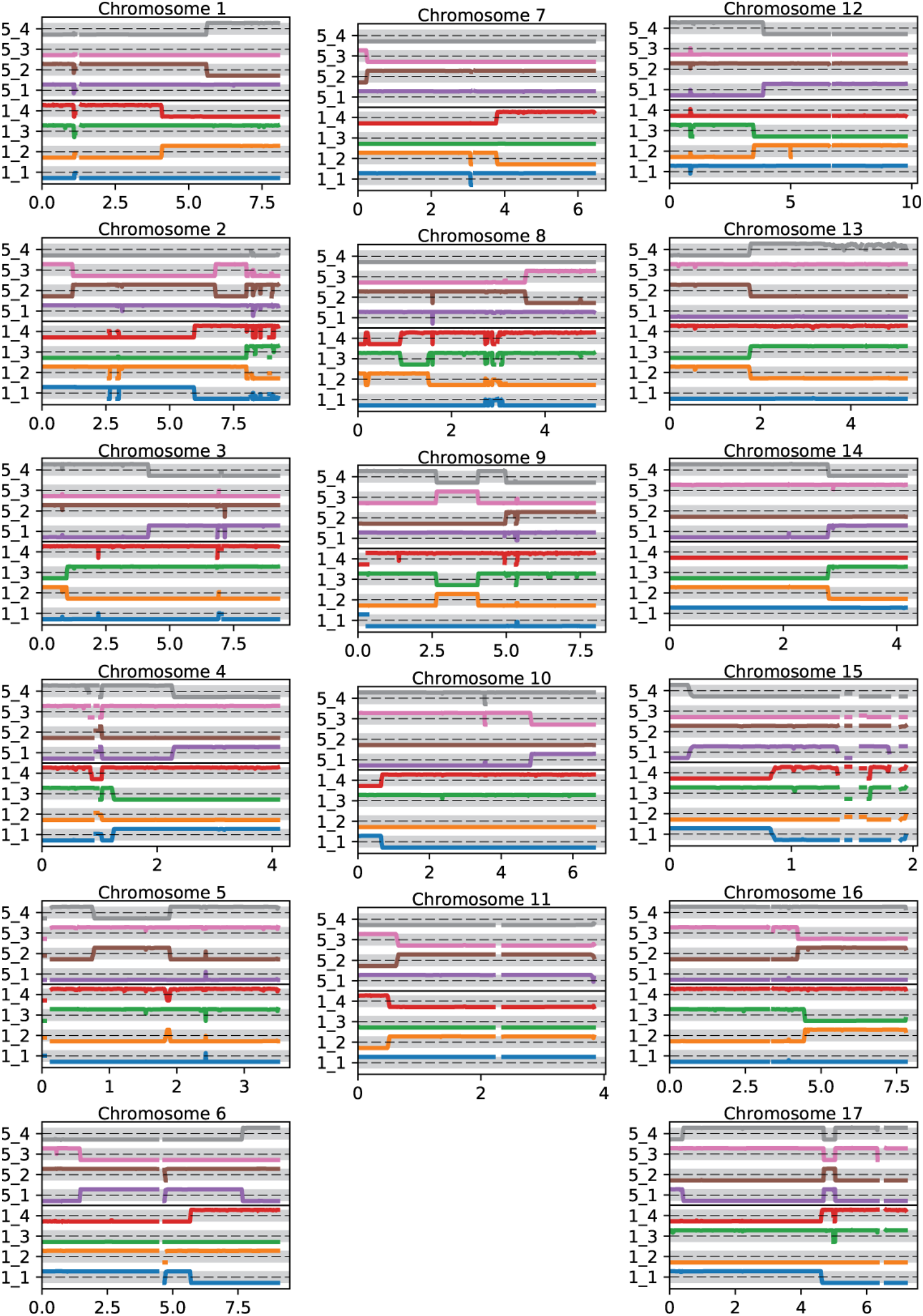
Genomic maps of the daughter cells resulting from two independent meiotic events are shown. Daughters 1_1 through 1_4 are from one meiotic event and 5_1 through 5_4 are from the second. The allele frequency is relative to CC2343 and the range of each shaded horizontal bar in is from 0 to 1, the dashed horizontal lines represent an allele frequency of 0.5.

**Figure S8.**
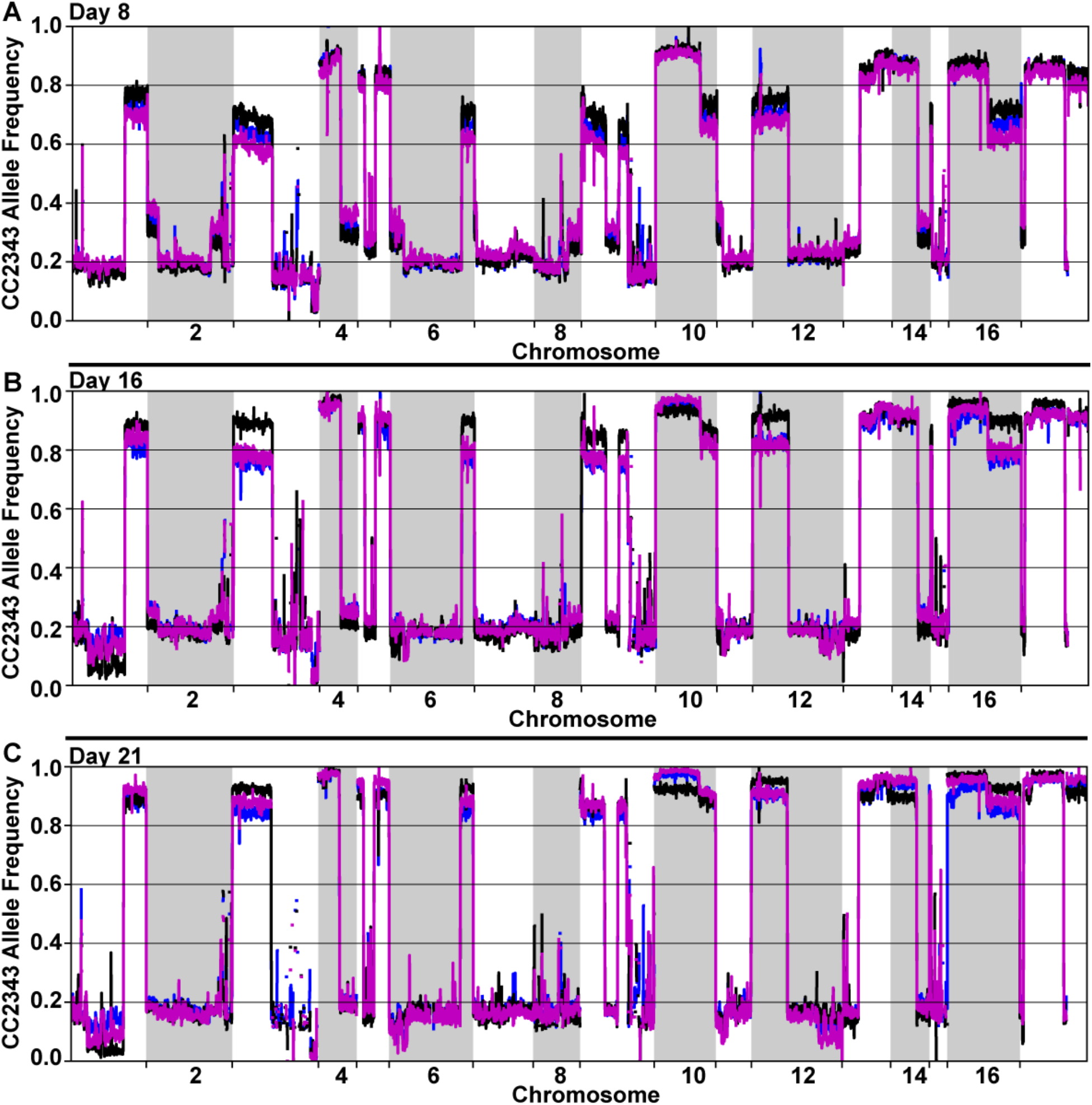
The allele frequency relative to CC2343 for individual F2 BC populations at days 8 (A), 16 (B), and 21 (C) are shown. Allele frequencies were averaged over 40 Kb windows sliding every 8 Kb. Line colors represent individual replicates.

**Figure S9.**
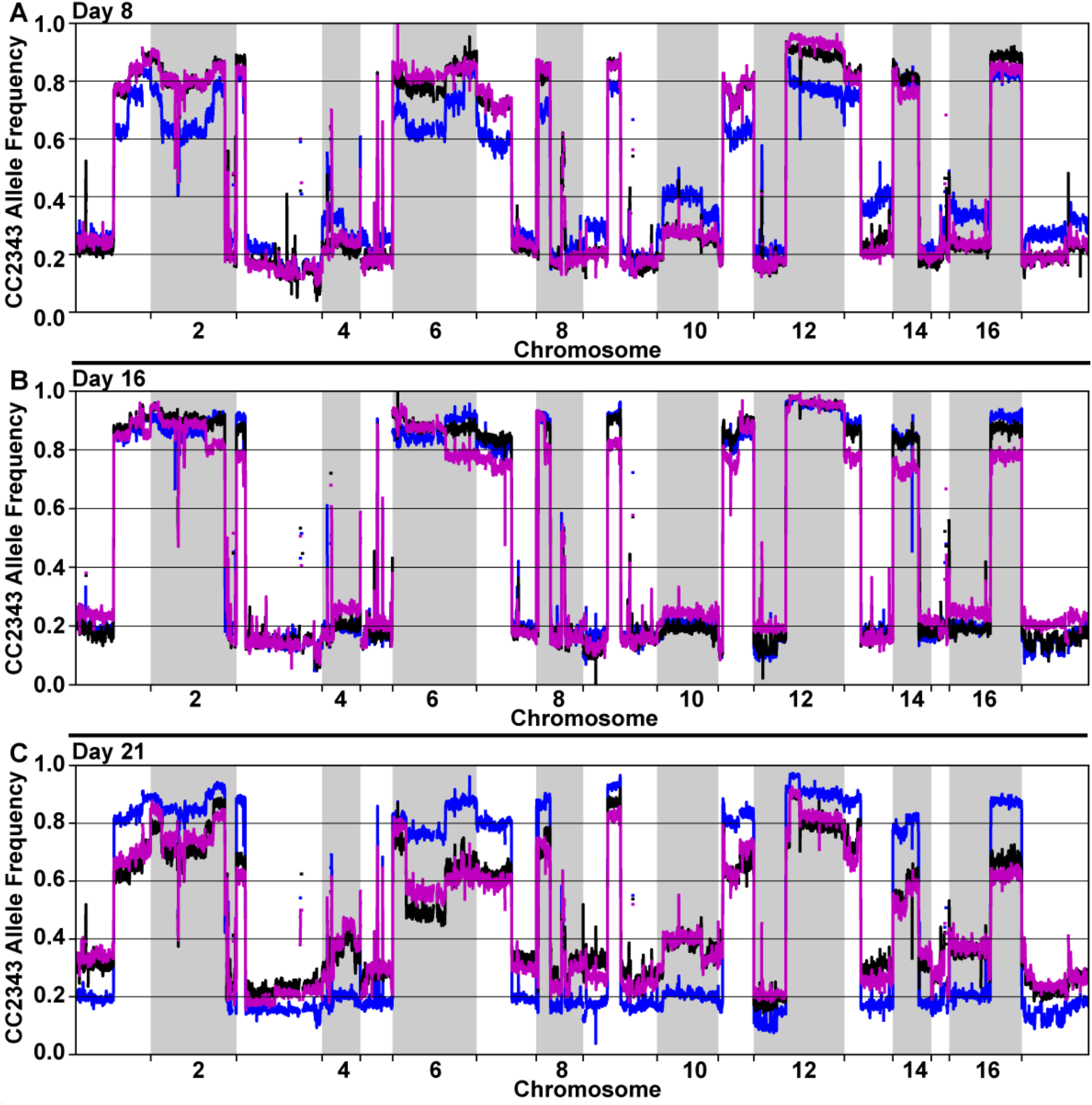
The allele frequency relative to CC2343 for individual F2 HO populations at days 8 (A), 16 (B), and 21 (C). Allele frequencies were averaged over 40 Kb windows sliding every 8 Kb. Line colors represent individual replicates.

**Figure S10.**
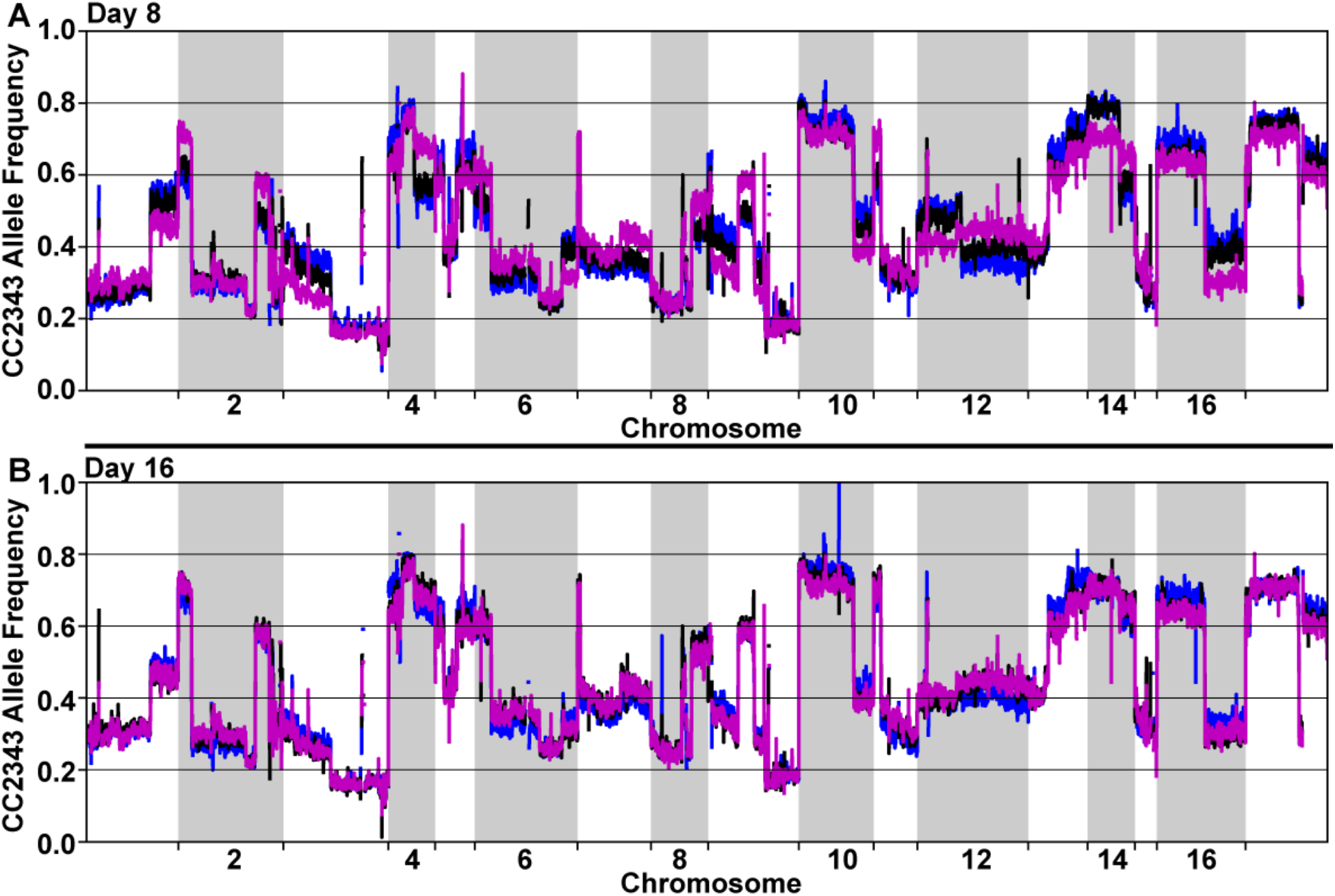
The allele frequency relative to CC2343 for individual F2 LS populations at days 8 (A) and 16 (B) are shown. Allele frequencies were averaged over 40 Kb windows sliding every 8 Kb. Line colors represent individual replicates.

**Figure S11.**
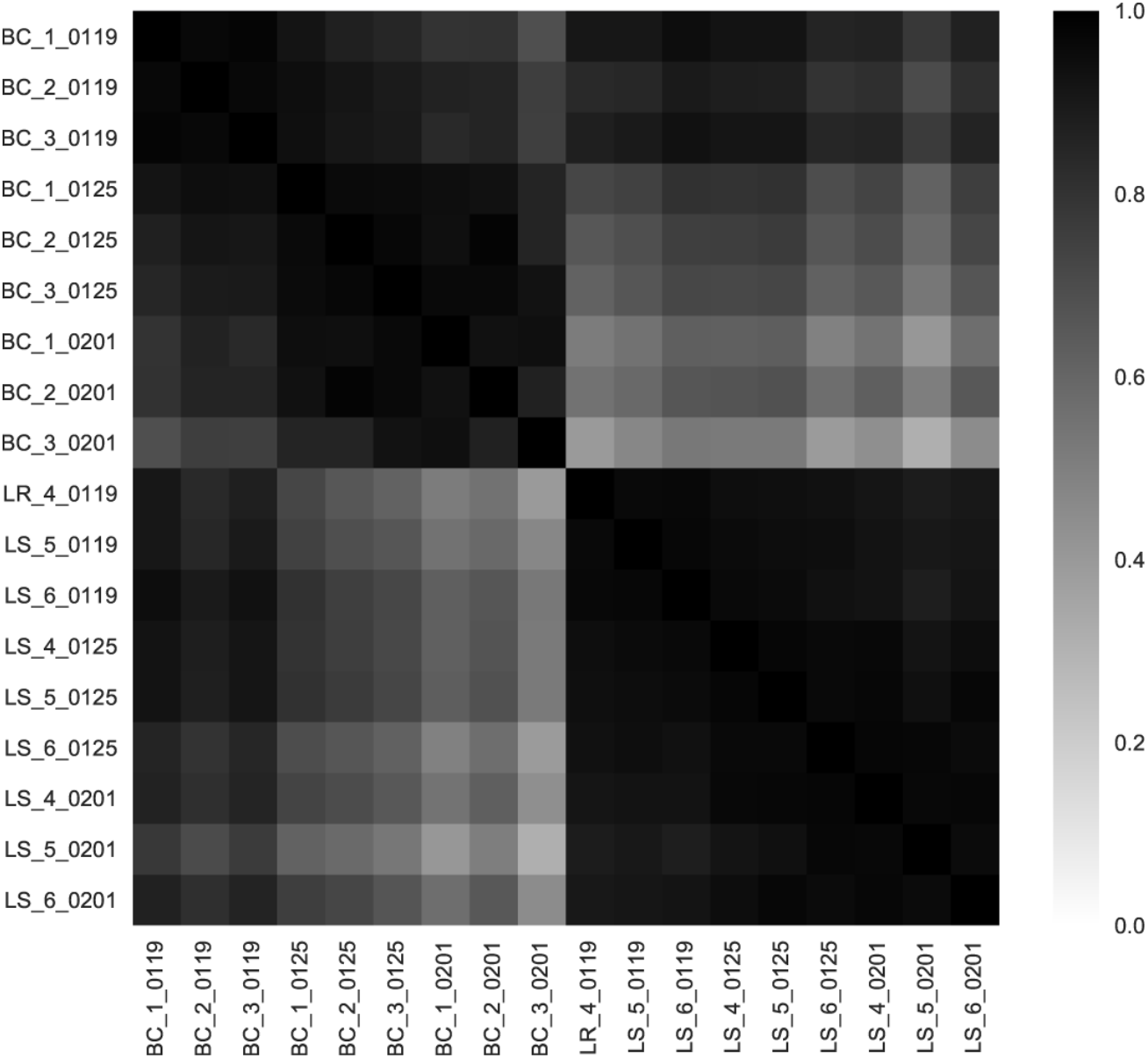
A high degree of reproducibility is shown using a matrix of Pierson Correlation Coefficients between the allele frequencies of replicate F2 populations exposed to the same environmental selection.

**Figure S12.**
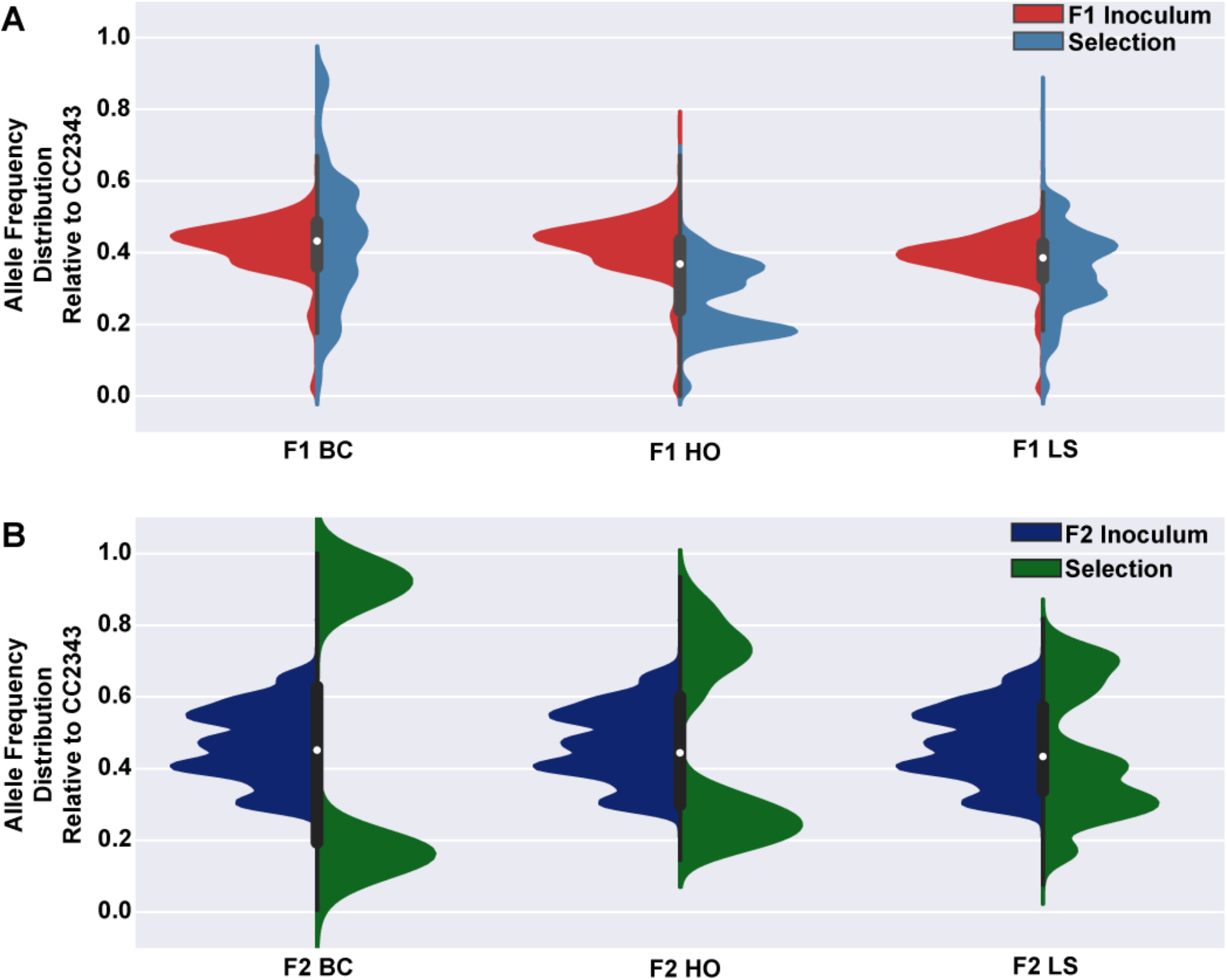
Allele frequency distributions in the F1 and F2 populations are highly divergent. (A) shows the allele frequency distribution of the inoculum (red curves) and the F1 competition populations (light blue curves). The distribution of the F1 BC and HO populations are after 31 days of culture while the F1 LS population was after 19 days. (B) Shows the allele frequency distribution of the F2 inoculum (dark blue curves) and the F2 populations (green curves) after 21 days of culture for the F2 BC and HO populations and 16 days for the F2 LS populations.

**Figure S13.**
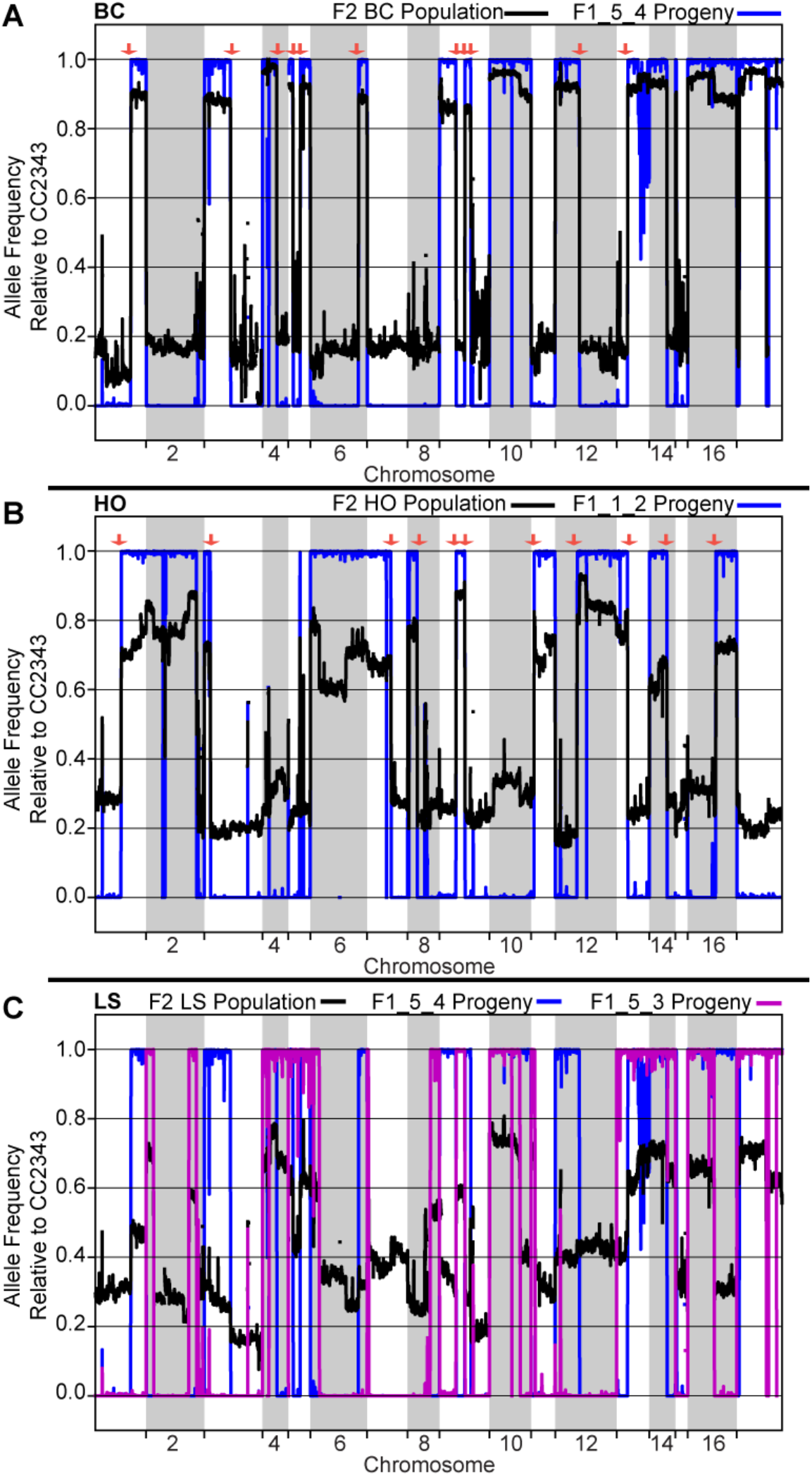
The survival of F2 progeny is heavily influenced by the genotype of the F1 parental. (A) The allele frequency distribution of the F2_BC population (black line) after 21 days of culture and the F1_5_4 meiotic progeny is shown. (B) Shows the allele frequency distribution of the F2_HO population (black line) after 21 days of culture, and the F1_1_2 meiotic progeny. (C) Shows the allele frequency distribution of the F2_LS (black line) after 16 days of culture and the F1_5_4 and F1_5_3 (blue and magenta lines, respectively).

**Figure S14.**
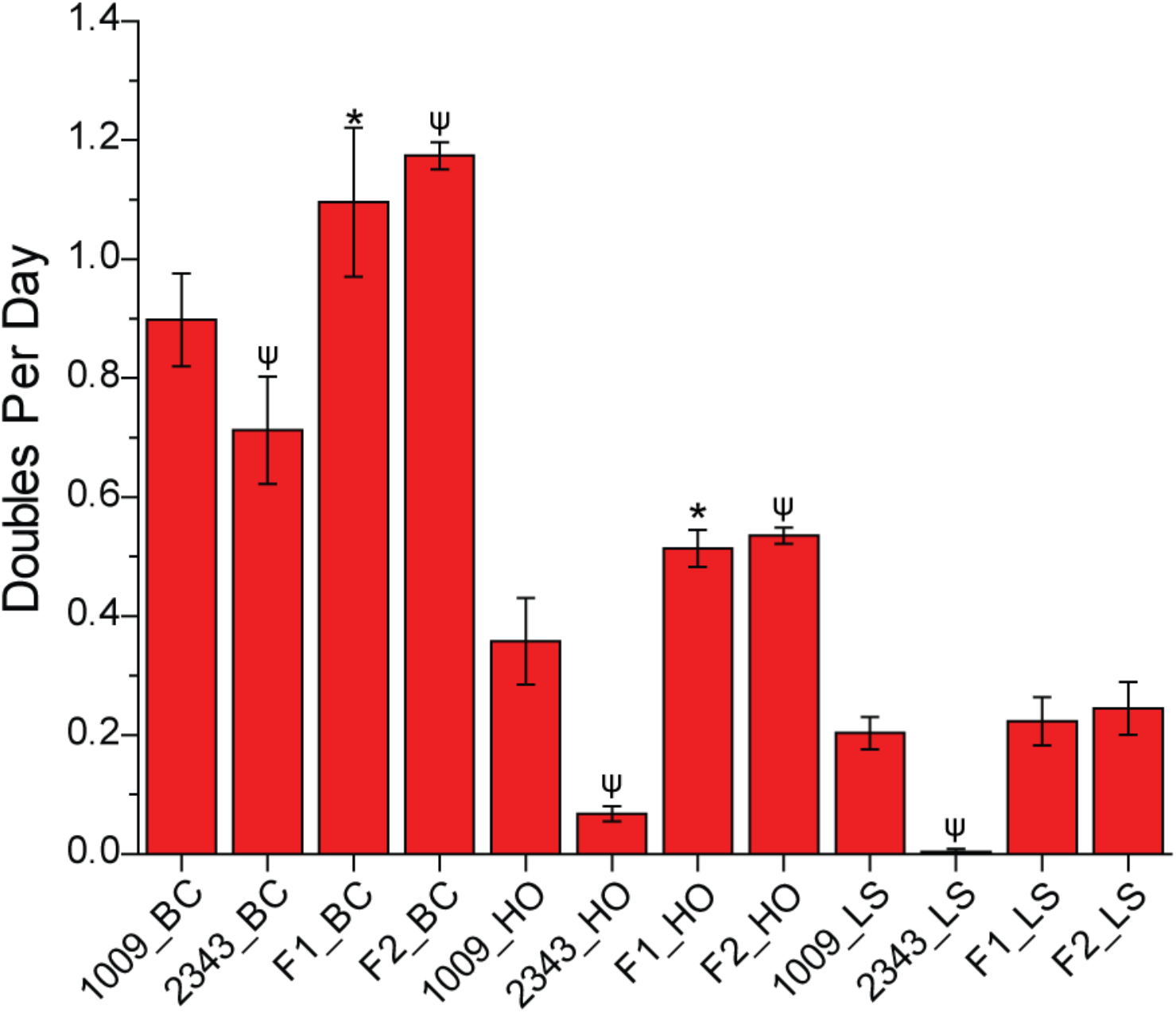
Populations showed increased doubling rates compared to parental lines. Doubles per day for F1 and F2 BC and HO populations were averaged from the last 5 full days of culturing. Error bars represent the standard deviation between biological replicates where, from left to right n = 8, 8, 5, 5, 5, 5, 3, 3, 5, 5, 3, 3. For the statistical analysis a two tailed t-distribution test was used and an * indicates a p value less than 0.01 and ψ represents a p value < 0.0005 for differences in doubles per day the each sample and CC1009 under the same conditions.

**Figure S15.**
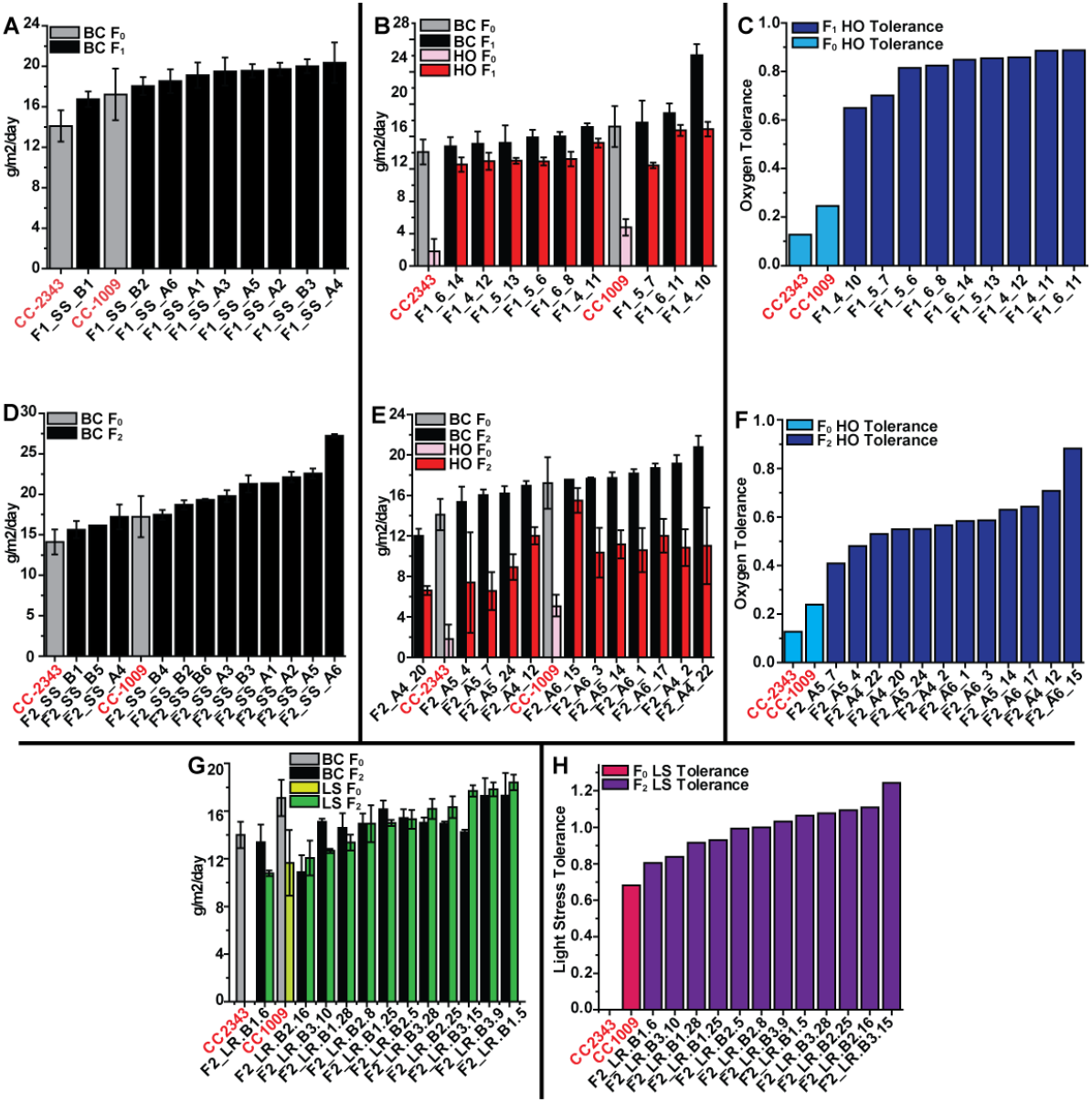
Meiotic progeny selected under polyculture conditions are enriched with strains with increased fitness. (A) Shows the daily productivity (in g of ash free dry weight produced per m^2^ of incident light) of the progenitor lines (F_0_) and F_1_ survivors isolated after 30 days of polyculture under BC conditions. (B) Shows the average daily productivity under BC conditions and HO conditions of the F_0_ and F_1_ survivors isolated after 30 days of polyculture under HO conditions. (C) Shows the oxygen tolerance of the F_0_ lines and the F_1_ HO survivors in panel B. (D) Summarizes a screen of the F_0_ lines and F_2_ progeny isolated after 21 days of polyculture under BC conditions. (E) Shows the average daily productivity under BC conditions and HO conditions of the parental F_0_ lines and F_2_ HO progeny isolated after 21 days of polyculture. (F) Shows the oxygen tolerance of the F_0_ parental lines and the F2_HO survivors in panel E. (G) Shows the average daily productivity under BC conditions and LS conditions of the parental F_0_ lines and F_2_ progeny isolated after 16 days of polyculture under LS conditions. (H) Summarizes the light stress tolerance of the lines shown in panel G. For the F_1_ and F_2_ progeny error bars represent the standard deviation between at least 3 daily growth values. For the F_0_ parental lines the error bars represents the standard deviation of the average daily productivity values between at 5 biological replicates for HO and LS conditions and 8 biological replicates for BC conditions.

**Figure S16.**
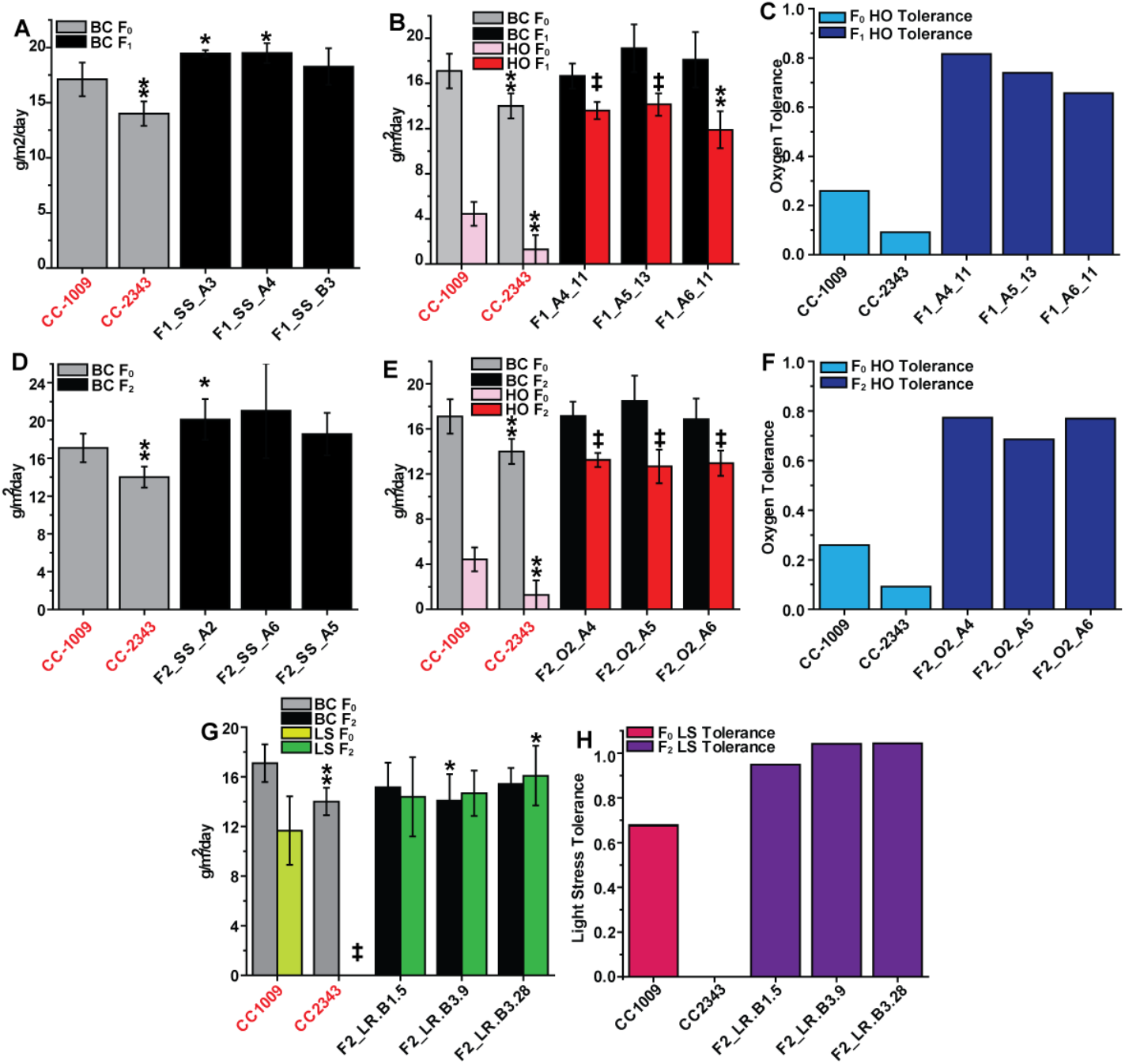
Strong heterosis persists in lines through multiple biological replicates. Panel A shows the daily productivity (in g of ash free dry weight produced per m^2^ of incident light) of the progenitor lines (grey bars) and choice F1 meiotic progeny (black bars) isolated after 30 days of polyculture under BC conditions. From left to right n = 8, 8, 3, 3, 3. Panel B shows the average daily productivity under BC conditions (grey and black bars) and HO conditions (pink and light red bars) of the parental and choice F1 progeny (respectively) isolated after 30 days of polyculture under HO conditions. From left to right n = 8, 5, 8, 5, 4, 4, 4, 3, 4, 3. Panel C shows the oxygen tolerance of the parental lines (dark blue bars) and the selected F1_HO survivors (dark grey bars) in panel B. Panel D shows the productivity of the progenitor lines (grey bars) and selected F2_BC survivors (black bars) isolated after 21 days of polyculture under BC conditions. From left to right n = 8, 8, 4, 4, 4. Panel E shows the average daily productivity under BC conditions (grey and black bars) and HO conditions (pink and red) of the parental and selected F2_HO progeny (respectively) isolated after 21 days of polyculture. From left to right n = 8, 5, 8, 5, 4, 4, 4, 4, 4, 3. Panel F shows the oxygen tolerance of the parental lines (light blue bars) and the selected F2_HO survivors shown (dark blue bars) in panel E. Panel G shows the average daily productivity under BC conditions (grey and black bars) and LS conditions (lime and green bars) of the parental and chosen F2_LS progeny (respectively) isolated after 16 days of polyculture. From left to right n = 8, 5, 8, 5, 4, 4, 4, 4, 4, 4. Panel H summarizes the light stress tolerance of the lines shown in panel G, pink bars represent the progenitor lines and purple bars represent F2_LS survivors. Error bars represent standard deviation of the averaged daily growth for each biological replicate. Asterisks denotes a maximum p-value of 0.05 from a two tailed t-distribution test while double asterisks denotes a maximum p-value of 0.005, ‡ represent a maximum p-value of 2e^-5^.

**Figure S17.**
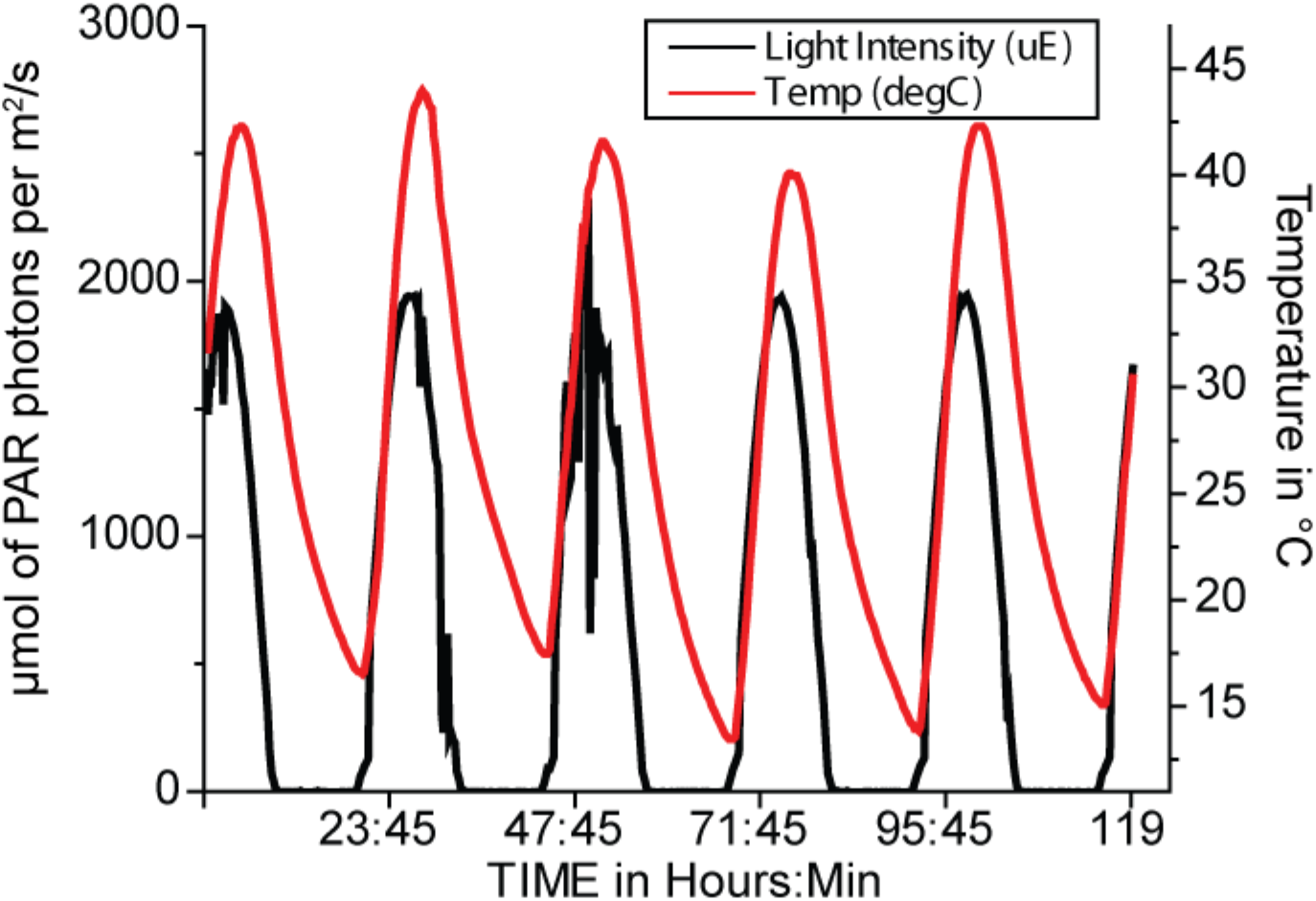
The light intensity (black line) and temperature (red line) during the environmental simulation selection is shown.

**Table S1.** Summarizes the genome coverage for each deep sequencing sample. Samples in red denote poor sequence alignments to the reference genome and failed the SNP calling pipeline.

| Experiment | Sample |  | day 9 | day 21 | day 25 | day 31 |
| --- | --- | --- | --- | --- | --- | --- |
| F1 HO | F1 Inoculum 1 | 42 |  |  |  |  |
|  | BC 1 |  | 40 | 38 | 42 | 42 |
|  | BC 2 |  |  |  |  |  |
|  | BC 3 |  | 41 | 42 | 43 | 44 |
|  | HO 1 |  | 41 | 43 | 42 | 43 |
|  | HO 2 |  | 44 | 41 | 40 | 43 |
|  | HO 3 |  | 42 | 42 | 43 | 56 |
|  |  |  | day 6 | day 12 | day 19 |  |
| F1 LS | F1 Inoculum 2 | 55 |  |  |  |  |
|  | BC 1 |  | 60 | 56 | 54 |  |
|  | BC 2 |  | 51 | 52 | 50 |  |
|  | BC 3 |  | 47 | 52 | 47 |  |
|  | LS 1 |  | 58 | 53 | 58 |  |
|  | LS 2 |  | 54 | 51 | 51 |  |
|  | LS 3 |  | 54 | 53 | 49 |  |
|  |  |  | day 8 | day 16 | day 21 |  |
| F2 | F2 Inoculum | 38 |  |  |  |  |
|  | BC 1 |  | 36 | 29 | 35 |  |
|  | BC 2 |  | 31 | 29 | 34 |  |
|  | BC 3 |  | 32 | 27 | 32 |  |
|  | HO 1 |  | 33 | 36 | 41 |  |
|  | HO 2 |  | 39 | 35 | 28 |  |
|  | HO 3 |  | 38 | 42 | 35 |  |
|  | LS 1 |  | 28 | 32 |  |  |
|  | LS 2 |  | 32 | 33 |  |  |
|  | LS 3 |  | 34 | 35 |  |  |
| Tetrad | F1_1_1 | 51 |  |  |  |  |
|  | F1_1_2 | 38 |  |  |  |  |
|  | F1_1_3 | 109 |  |  |  |  |
|  | F1_1_4 | 52 |  |  |  |  |
|  | F1_5_1 | 56 |  |  |  |  |
|  | F1_5_2 | 35 |  |  |  |  |
|  | F1_5_3 | 41 |  |  |  |  |
|  | F1_5_4 | 92 |  |  |  |  |
| Parental | CC1009 | 44 |  |  |  |  |
|  | CC2343 | 62 |  |  |  |  |

**Table S2.** The change in Allele Frequency (ΔAF) from the Inoculum Relative to CC1009, and the number of generations per population

| Experiment Condition |  | day 9 |  | day 21 |  | day 26 |  | day 33 |  |
| --- | --- | --- | --- | --- | --- | --- | --- | --- | --- |
| | | ΔAF | Generations | ΔAF | Generations | ΔAF | Generations | ΔAF | Generations |
| F1 HO | BC | 7.2e-4 | 9.3 ± 1.38 | 5.4e-3 ± 2.0e-2 | 21 ± 4.6 | 8.9e-3 ± 1.4e-2 | 26 ± 4.3 | 2.3e-2 ± 5.9e-3 | 33 ± 5.6 |
| | HO | 1.3e-3 ± 2.3e-3 | 4.5 ± 0.57 | 0.12 ± 1.7e-2 | 9.1 ± 1.5 | 0.11 ± 1.4e-2 | 11 ± 1.6 | 0.15 ± 4.9e-2 | 14 ± 1.8 |
|  |  | day 6 |  | day 12 |  | day 19 |  |  |  |
| F1 LS | BC | 9.6e-3 ± 3.1e-2 | 6.0 ± 0.68 | 4.7e-3 ± 9.1e-2 | 12 ± 1.4 | -3.1e-2 ± 6.1e-3 | 19 ± 2.4 | | |
| | LS | 2.1e-2 ± 4.8e-4 | 3.0 ± 0.38 | 4.3e-2 ± 4.7e-4 | 3.9 ± 0.51 | 2.8e-2 ± 4.9e-3 | 5.7 ± 0.83 | | |
|  |  | day 11 |  | day 19 |  | day 25 |  |  |  |
| F2 All | BC | 2.2e-4 ± 0.005 | 13 ± 0.59 | -3.2e-2 ± 8.3e-3 | 23 ± 0.91 | -4.6e-2 ± 7.1e-3 | 29 ± 0.92 | | |
| | O2 | -1.0e-2 ± 7.1e-3 | 5.2 ± 0.26 | -1.1e-2 ± 2.8e-3 | 8.6 ± 7.0e-2 | 4.4e-3 ± 3.4e-3 | 12 ± 0.22 | | |
| | LR | 2.2e-2 ± 3.6e-3 | 5.9 ± 0.38 | 2.3e-2 ± 1.5e-3 | 8.2 ± 1.0 | | | | |

## Materials and Methods

### Strains, Media and Generation of Progeny

*Chlamydomonas* strains CC1009 (mt-) and CC2343 (mt+) were obtained through the Chlamydomonas Resource Center. CC1009 and CC2343 were crossed using methods described by Jiang and Stern (80) (2009). We identified 204 F1 mt- progeny using a colony PCR screen and mating type specific primers (MTA and MTD) as described in Lin and Dutcher(56) (2015). The 246 F2 progeny population was generated by dissecting two F1 zygotes and crossing the reciprocal mating types of each tetrad (each mt- with mt+ from each tetrad as shown in Fig. 3A) for total of 8 F1 crosses (∼30 lines from each cross). Cultures were maintained on either Sueoka’s high salt media(81)or 2NBH media, which is a Bristol media with 2 x sodium nitrate and Hutner’s trace elements added(82).

For the zygote selection experiments. F2 strains were crossed and zygospores were harvested from plates and pooled prior to hatching. For the pond like conditions population, pooled zygotes were hatched in 2NBH media under low light (250 µmol of photosynthetically active radiation photons per m^2^/s) with gentle stirring in the ePBR reactors with a column height of ∼5cm. Hatching zygospores for the high salt selection experiments was the same except the population was hatched in 2NBH media containing 20 g/L of Instant Ocean Salts.

### Growth and Competition Conditions

To achieve highly controlled and reproducible conditions, we used a set of environmental photobioreactors (ePBRs)(67). Cultures were maintained at a constant density by automatic dilution with 5ml of fresh media if the turbidity was measured above the set point, until the turbidity decreased below the set point. In this way, the relative biomass growth for the cultures over a time range could be estimated by the number of dilutions, as describe in the following section (see also Lucker and Hall et al. 2014). For competition experiments the chlorophyll concentration was maintained between 4 and 5 µg chlorophyll ml^-1^. The ePBR culture height was set to 15 cm using a volume 330 ml of 2NBH media. For individual phenotyping conditions, cultures were pre-conditioned to grown in ePBRs to a chlorophyll 4 µg ml^-1^ and maintained in turbidistat mode using the standard light conditions for at least 3 days prior to measuring productivity. For the light stress (LS) and high oxygen (HO) competition experiments, the pre-conditioning time was reduced to a single day to avoid imposing long-term selection under the baseline conditions (BC). For the BC and HO conditions, standard illumination was provided on a 14:10 hour (light:dark) diurnal cycle simulating a cloudless day, with light intensity ascending to a zenith with maximum photosynthetically active radiation (PAR) of 2000 µmol photons m^-2^ s^-1^, and descending until dark, delivered in a sinusoidal form, as illustrated in Figure S1. For the LS regime, the standard illumination days were alternated with a series of three “light starvation” days, which consisted of a simple, 14:10 hour rectangular wave with a PAR intensity of 50 µmol photons m^-2^ s^-1^, Figure S1. All cultures were stirred at 200 rpm using a 28.6 mm by 8 mm Teflon coated stir bar. Gas for BC and LS conditions was 5% CO_2_ in air and gas for HO was 5% CO_2_ in O_2_. Gas delivered through a 5 mm gas dispersion stone with a porosity of 10-20 microns at a flow rate of 350 ml / min for 60 seconds every hour. Culture temperatures were maintained at RT for the F1 and F2 competition and 25 ^○^C for monoculture phenotyping of parental lines and competition survivors.

The ePBR vessel, stirring and gas delivery was the same as the BC and LS experiments described above. The temperature and light intensity for the for the pond like environmental simulation is shown in figure S17. For the high salt selections and screening the ePBR configuration was the same as the BC conditions except the media contained 20 g/L of Instant Ocean Salts and the temperature was maintained at 28 ^○^C. It should be noted that the light for the ePBR for both the pond like simulation and high salt selection is the next generation ePBR light and is a 50 W white LED with an optical columnator.

### Biomass Productivity

Biomass productivity was determined by multiplying the number of daily turbidistat dilutions (5 ml per dilution) and the Ash Free Dry Weight (AFDW) then dividing by the area of the top of the ePBR culture vessel at 15 cm (0.002687 m^2^). Ash free dry weight was determined by concentrating 35 ml of culture onto a Whatman CF/F glass filter and dried over night at 104 ^○^C prior to weighing for the “dry weight”. Organic matter was removed from the filter by combusting the samples at 550 ^○^C for a minimum of 30 minutes prior to weighing the sample for the “ash weight”. The AFDW is the dry weight minus the ash weight.

### Genome Sequencing

DNA from all samples were isolated described in Fawley and Fawley (2004). Genomic DNA library generation was performed by the Michigan State University Genomics Core Facility using the Illumina TruSeq Nano DNA Library (www.illumina.com) with dual 8bp index adapters. Libraries were checked for quality and quantified using Qubit dsDNA HS, Caliper LabChipGX HS DNA (www.perkinelmer.com) and Kapa Biosystems Illumina Quantification qPCR assays (www.perkinelmer.com). Libraries pooled for multiplexed sequencing and loaded on 2 lanes of an Illumina HiSeq 2500 High Output flow cell (v4) and sequencing was performed with HiSeq SBS reagents (v4) in a 2×125bp paired end format. Base calling was done by Illumina Real Time Analysis (RTA) v1.18.64 and output of RTA was demultiplexed and converted to Fastq format with Illumina Bcl2fastq v1.8.4. Samples for the tetrad analysis, F2 competition and the parental lines were prepared similar to the previous samples, but sequenced and processed using an Illumina HiSeq 4000 flow cell in a 2×150bp paired end format using HiSeq 4000 SBS reagents. Base calling was done by Illumina Real Time Analysis (RTA) v2.7.6 and output of RTA was demultiplexed and converted to FastQ format with Illumina Bcl2fastq v2.19.0. The average genomic sequencing depth of the tetrad and F2 experiments is reported in Table S1. Genomic DNA read pairs were aligned the *Chlamydomonas* reference genome v5.0 (JGI v5.0 assembly, JGI annotation based on Augustus u11.6) using the bowtie2/2.2.3 aligner. For each sam file output, the file was converted to bam and the reads were sorted, read groups were replaced with a single group, mate pair information was fixed and duplicated mates were removed using picardTools/1.113 (https://github.com/broadinstitute/picard/). Reads were realigned to the reference genome using GATK3.1.1(83). Variant bases were called using SamTools/0.0.19(84) and output was filtered for repeats and formatted into the variant call format (VCF) using vcftools/0.1.12a(85).

### Identification of quantitative singleton single nucleotide polymorphisms (SNPs)

To map the allele frequencies of CC2343 and CC1009 loci in polyculture populations of meiotic progeny we required a list of single nucleotide polymorphisms (SNPs) from the parental strains that were reliably quantitative. Using Illumina reads from CC2343 and CC1009 we generated computational populations by pooling reads into 3 sets of 24 million reads containing either 75%, 50%, or25% of CC2343 and CC1009. After aligning each computational population we determined the allele frequency for each SNP site in the VCF output. The SNP sites for each population were rough filtered with the ∼2.6 million SNPs between CC2343 and CC1009 relative to the sequenced CC503 strain (71). Sites assigned to CC2343 and CC1009 were deemed quantitative if they deviated less than 15% from the known target frequency of the computational population (filtering > 1 million SNPs), resulting in over 1.6 million SNPs assigned to CC2343 (∼ 1.56e^6^ SNPs) or CC1009 (∼1.07e^5^ SNPs).

### Allele Frequency determination and Identification of selection enriched genomic loci (SEGLs)

For each population sequenced the percentage of variant vs reference calls was determined from the output VCF file prior to assigning any site as either progenitor, CC1009 or CC2343 using the quantitative SNP list described above. The SNP frequency information for both parents was then merged relative to the CC2343 parental line. To reduce the influence of local allele frequency fluctuations, averaged allele frequencies for genomic regions were determined using 40 Kb windows and a step size of 8 Kb down each chromosome.

We used the allele frequency output to map Selection Enriched Genomic Loci (SEGL) regions with significant changes in allele frequency from the inoculum for each F1 population and environmental condition (BC, HO or LS) with two or more replicates. First, we calculated the allele frequency differences between the two F1 inoculum samples, which formed a distribution that was approximately normal. After taking the absolute values of the differences, a folded-normal distribution model was fit to the absolute difference distribution with the foldnorm function in SciPy(86). We calculated the differences in allele frequencies between each environmental condition/time point and its respective inoculum for each window. The allele frequency difference for each window was used to determine the probability (p) of observing a difference of that magnitude or greater using the cumulative distribution function of our folded-normal model. Given the overlap between adjacent windows and the effects of linkage disequilibrium, these p-values are likely not-independent, therefore we could not apply the standard multiple-testing procedure and decided to be conservative in our estimates of truly significant changes by using a stringent cutoff (p<1e-14). We define SEGLs, for each condition/timepoint, as consecutive windows with p<1e-14 for all replicate samples merged into a contiguous region ≥60Kb.

